# Interplay between ferroptosis and guttae in an early-onset murine model of Fuchs’ endothelial corneal dystrophy (FECD)

**DOI:** 10.64898/2026.07.09.737597

**Authors:** Karin W. Handel, Jaegook Lim, Hiroko Iwashita, Sabina Khan, Hanna Shevalye, Sangwan Park, Nayeli Echeverria, Michelle Ferneding, Meher J. Khan, Karolina P. Roszak, Gabriella L. Donovan, Mako Iwamoto, Jaeho Shim, Laura J. Young, Monica Ardon, Sophie Le, Brian C. Leonard, Jessica M. Skeie, Mark A. Greiner, Sara M. Thomasy

## Abstract

*Col8a2^Q455K/Q455K^* (Q455K) mice exhibit features of early-onset Fuchs’ endothelial corneal dystrophy (FECD), including decreased endothelial cell density (ECD) and guttae formation. Within the context of these clinical features, this study longitudinally evaluates ferroptosis in Q455K and wild-type (WT) mice using *in vivo* imaging, PCR and immunohistochemistry. Fifty-six Q455K and 56 WT mice were evaluated from 3 to 24 months of age with in vivo confocal microscopy; ECD and guttae were measured. Ferroptosis marker expression was determined with PCR and immunohistochemistry (IHC). Data were analyzed using two-way ANOVA with Tukey’s post hoc test, Chi-square test and a paired t-test. The ECD significantly decreased in both groups from 3 to 24 months of age, but more markedly in Q455K (2285 ± 317 to 1012 ± 58 cells/mm²) versus WT mice (2714 ± 139 to 2057 ± 149 cells/mm², *P*<0.0001). Guttae were observed exclusively in Q455K mice beginning at 3 months of age and increased over time (*P*=0.0003). The Q455K mice demonstrate guttae at the vertices of corneal endothelial cells rather than their centers (74.3% vs. 25.7%*P<*0.001). Expression of ferroptosis-related genes (*Tfrc, Slc40a1, Ftl1, Gpx4*) were significantly increased in the Q455K versus WT mice (*P*<0.05). Furthermore, corresponding protein expression (transferrin receptor 1, ferroportin, ferritin and glutathione peroxidase 4) was significantly elevated adjacent to guttae in Q455K versus WT mice (*P*<0.05). These findings implicate guttae in the initiation of ferroptosis as it relates to the pathophysiology of FECD and provide an optimal window for testing novel FECD therapies using this murine model, particularly those that target ferroptosis.

**Significance Statement:** Fuchs’ endothelial corneal dystrophy (FECD) is a major cause of corneal blindness, yet the mechanisms driving endothelial cell loss remain unclear. Using an early-onset Q455K murine model, we show that disease onset occurs by 3 months of age and progresses with characteristic corneal guttae, endothelial cell loss, and morphological abnormalities replicating human FECD. By identifying ferroptosis adjacent to guttae in the Q455K, we demonstrate this mechanism is not specific to genetic mutations, but rather a conserved driver of corneal endothelial loss within the context of guttae formation. These findings establish an FECD progression timeline and reveal a mechanistic link between ferroptosis and guttae, validating this model as a critical tool for evaluating targeted therapeutic strategies.

## Introduction

Fuchs’ endothelial corneal dystrophy (FECD) is a progressive degenerative disease of the corneal endothelium that is marked by the gradual loss of corneal endothelial cells (CEnC), thickening of Descemet’s membrane (DM), and the formation of posterior excrescences within the DM, termed guttae (1–6). These cells maintain corneal transparency by regulating stromal hydration via sodium-potassium ATPase (Na⁺/K⁺-ATPase) pumps that facilitate fluid efflux out of the cornea and tight junctions maintained by zonula occludens-1 (ZO-1) that form a barrier to prevent aqueous humor entry into the cornea. Due to their limited proliferative capacity *in vivo*, progressive loss of CEnCs results in endothelial dysfunction and subsequent corneal edema (5, 7).

Fuchs’ endothelial corneal dystrophy is classified into 2 forms: late-onset FECD, typically manifesting clinical signs around the age of 40 years, and early-onset FECD, where findings may appear before the age of 10 (1, 8, 9). Two separate autosomal dominant point mutations. L450W (1) and Q455K (9) in the *COL8A2* gene, which encodes the α2 chain of type VIII collagen, cause early-onset FECD. Murine models carrying these mutations were developed, with Q455K mice demonstrating a more severe phenotype compared to L450W mice (1, 3, 7–12). However, studies detailing the onset and clinical progression of disease in these models are lacking.

The progression of FECD is closely linked to endothelial dysfunction and is driven by biophysical disturbances that arise from structural alterations in DM, including the accumulation of guttae and changes in extracellular matrix (ECM) composition. Methot and colleagues demonstrated in human corneas that CEnCs adjacent to guttae exhibit reduced pump activity and diminished mitochondrial volume, clearly indicating that guttae negatively affect adjacent CEnCs (13). In murine FECD models, just as in human patients with this condition, the presence, number, and distribution of guttae serve as critical morphological indicators of disease severity and progression, providing a valuable marker for evaluating disease progression and response to therapeutic interventions (3, 14–16). However, detailed quantification of guttae and how they drive CEnC death is understudied (17).

A key driver of FECD pathogenesis is compromised cellular defense against oxidative stress (18, 19). In FECD, CEnCs exhibit increased sensitivity to oxidative damage, with several factors contributing to increased vulnerability to lipid peroxidation. Adequate amounts of glutathione (GSH) and proper activity of glutathione peroxidase 4 (GPX4), an enzyme that reduces lipid peroxides in a GSH-dependent manner, are critical in preventing iron-dependent lipid peroxidation and ferroptosis, a form of nonapoptotic oxidative cell death. Patients with FECD demonstrate reduced GPX4 versus controls, indicating enhanced susceptibility to ferroptosis, an iron-dependent cell death mechanism (18, 20–22). Although the mechanism of ferroptosis in FECD and its potential for therapeutic targeting have been described in humans with TCF4 disease, ferroptosis has not been investigated in animal models nor other FECD genetic models, as a mechanism underlying the disease’s increased oxidative sensitivity (18, 23). Given these findings, the Q455K mouse model, already exhibiting phenotypic changes consistent with human FECD, could serve as a relevant *in vivo* model to explore the mechanistic role of ferroptosis in disease progression. Thus, the aims of this study were to quantify the onset and progression of corneal pathology over time in the Q455K mouse model, and to assess the involvement and extent of ferroptosis in the pathophysiology of this model and in human FECD patients.

## Results

### Progressive endothelial cell loss with aberrant morphology occurs in Q455K mice without corneal edema

When comparing all age groups, endothelial cell density (ECD) was significantly reduced in the Q455K versus wildtype (WT) mice (*P<*0.0001; **Figure 1A, 2**). An age-related decline in ECD was observed in both genotypes over time but was more pronounced in Q455K (2285±317 cells/mm² at 3 months and 1012±58 cells/mm² at 24 months) compared to the WT mice (2714±139 cells/mm² at 3 months to 2057±149 cells/mm² at 24 months). In both genotypes, ECD at 3 months of age was significantly greater than any other age group (*P≤*0.0002, **Supplementary Figure 1**). For both the WT and Q455K genotypes, no sex differences in ECD were identified (**Supplementary Figure 2**). Despite the profound reduction in ECD in the Q455K mice, none developed spontaneous corneal edema by 24-months of age and no differences in central corneal thickness (CCT) were identified between groups or over time (**Figures 1B, 2**). Na⁺/K⁺-ATPase is a critical transporter for maintaining corneal dehydration and was present in the periphery of the CEnC along the basolateral membranes in a lobulated pattern in WT mice of all ages as well as Q455K mice ≤6 months of age. However, the Q455K mice ≥9 months of age demonstrated increased Na⁺/K⁺-ATPase expression along the basolateral membrane particularly in the largest cells presumably to compensate for CEnC loss (**Figure 3A-B**).

**Figure 1.**
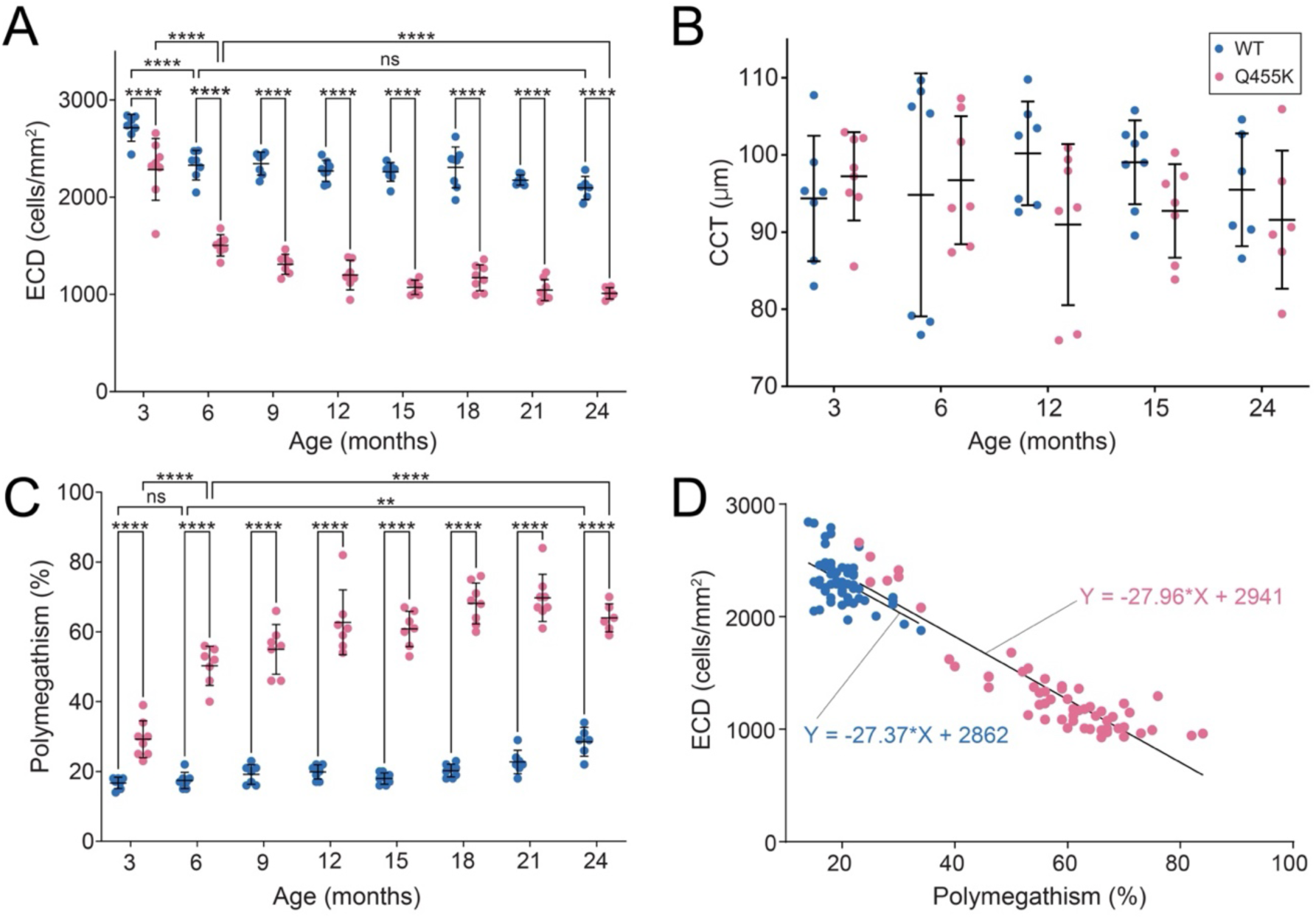
Endothelial cell density (ECD) was significantly decreased and polymegathism significantly increased in Q455K versus WT mice as well as over time in both groups; a strong, inverse correlation was observed between ECD and polymegathism. However, central corneal thickness (CCT) did not significantly differ between the two groups or over time. As shown in **(A)**, an age-related decline in ECD was observed in both genotypes over time but was more pronounced in Q455K (2285 ± 317 at 3 months and 1012 ± 58 cells/mm² at 24 months) versus WT mice (2714 ± 139 at 3 months to 2057 ± 149 cells/mm² at 24 months). In both genotypes, ECD was significantly higher at 3 months of age versus all other age groups studied (*P≤*0.0002, Supplementary Figure 1). However, CCT, as measured by FD-OCT, did not differ between the Q455K and WT ranging from 76-107 and 77-110 µm respectively **(B)**. Polymegathism was significantly increased in Q455K versus WT mice (*P<*0.0001; **C)**. At 3 months of age, polymegathism was already 29.3 ± 7.6 in the Q455K mice versus 16.7 ± 1.4% in the WTs (P<0.0001). Within the WT group, the 24-month age group had significantly greater polymegathism than all other age groups (*P<*0.05). Polymegathism was negatively correlated with ECD in both the WT and Q455K groups (*r*=−0.4378, *P=*0.0008 and *r*=−0.843, *P*<0.0001 respectively; **D).** \**P<*0.05, \*\**P<*0.01, \*\*\**P<*0.001, and \*\*\*\**P<*0.0001, ns is not significant.

**Figure 2.**
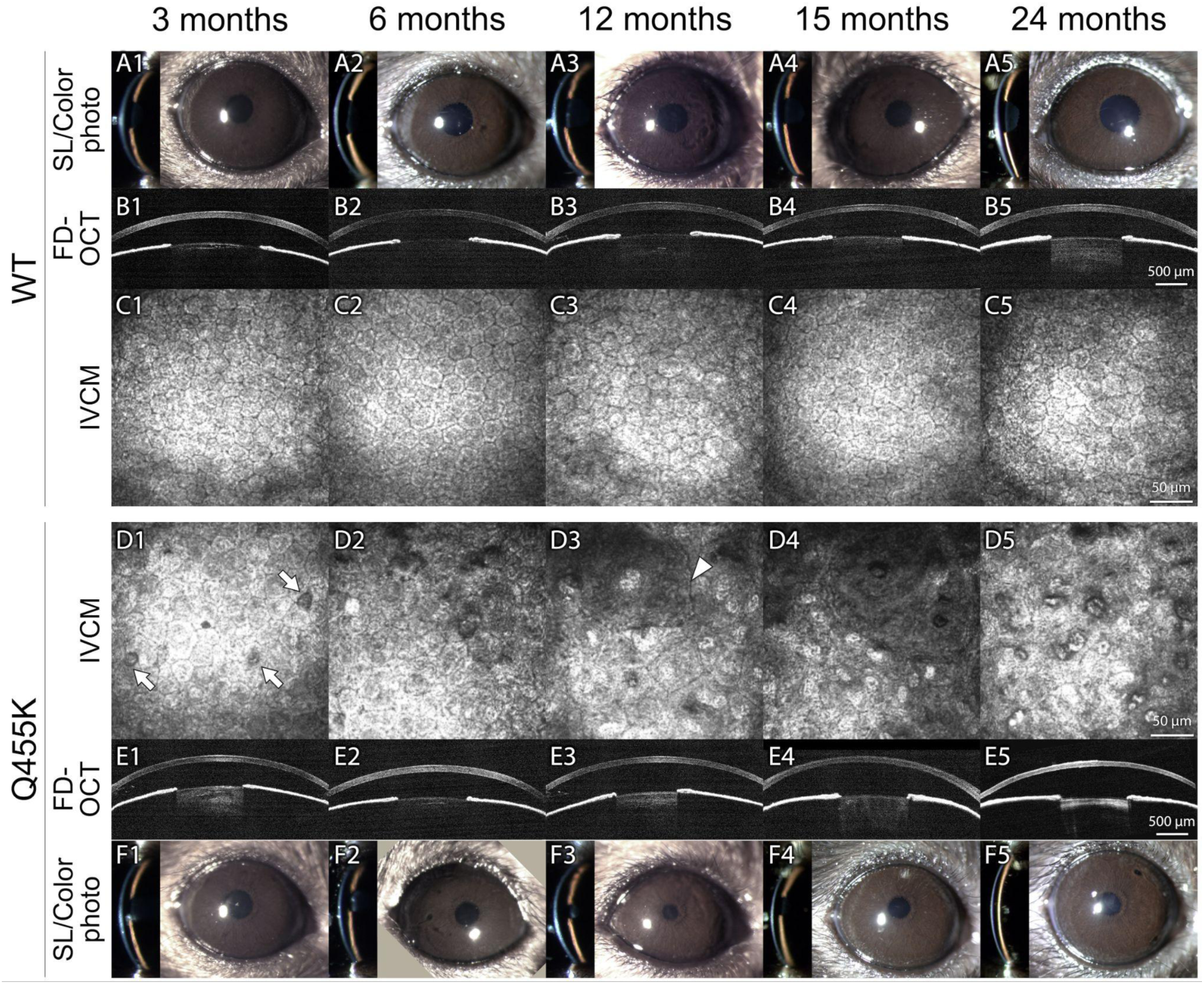
Corneal endothelial cell (CEnC) morphology dramatically differed between WT and Q455K mice but no difference in corneal thickness and clarity were observed between groups. The corneas of WT mice maintained normal clarity (**A1-A5**) and thickness (**B1-B5**) within all age groups. A representative 3-month WT mouse demonstrates normal CEnC density and morphology with a dense corneal endothelium (**C1**). At that age, the WT ECD is 2714 ± 139 cells/mm² and polymegathism is 17±1.3% (**C1**). With age, cell size gradually increases (polymegathism 16.7±1.4% at 6 months to 28.2±4% at 24 months) with mild changes in hexagonality (**C2-C5**). Within the Q455K group, a lower ECD (2285 ± 317 cells/mm²) and mild polymegathism (29.3% ± 7.6%) is identified at 3-months of age as exemplified in **D1**, A gradual decrease in ECD and increase in polymegathism is observed in the Q455K mice that culminates at 1012 ± 58 cells/mm² and 64%±8.6% by 24 months of age (**D1-D5**). The increase in polymegathism is notable already from 12 months of age (white arrowhead **D3**). Additionally, guttae are identified at 3 months of age in Q455K mice (**D1**, white arrows). With age, guttae density and guttae size/area gradually increase to 273 ± 37 guttae/mm² and 7.2±1.5 GAR by 24-months of age (**D1-D5**). In Q455K mice, no differences in CCT (**E1-E5**), or corneal clarity (**F1-F5**) were observed over time. Individual data for Q455K mice 1 (**D1-F1**), 15 (**D2-F2**), 23 (**D3-F3**), 31 (**D4-F4**), 55 (**D5-F5**) and for WT mice 6 (**A1-C1**), 10 (**A2-C2**), 23 (**A3-C3**), 37 (**A4-C4**) and 57 (**A5-C5**) can be found in supplementary tables 2 and 3 respectively.

**Figure 3.**
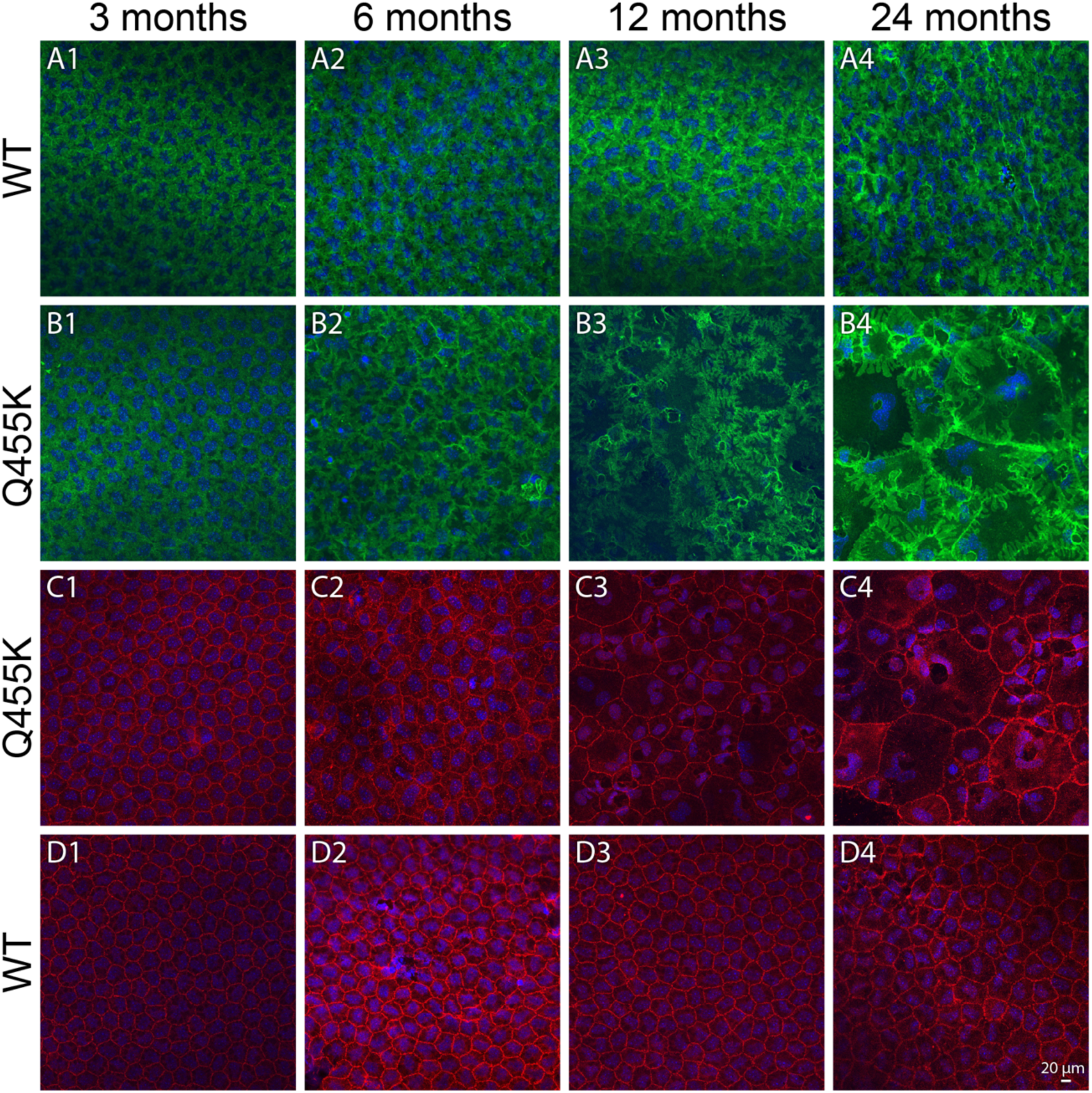
In Q455K CEnCs, increased basolateral expression of Na⁺/K⁺-ATPase (A-B, green) and polymegathism using zonula occludens-1 (ZO-1; C-D, red) is documented compared to WT CEnCs. Both WT and Q455K mice demonstrated discrete localization of Na⁺/K⁺-ATPase and ZO-1 to the cell periphery but they occupy basolateral and apical junctions, respectively. The Na⁺/K⁺-ATPase is observed along a widened basolateral junction in the Q455K mice that further increases in density at 12 months of age (**B3**). Using ZO-1, a loss of hexagonal morphology and an increase in cell size were evident in Q455K mice from 6 months of age, with worsening in cell structure by 12 months of age (**C3**). Guttae are denoted by a white asterisk (**C4**). Scale bar equivalent to 20 μm. Nuclei are stained with DAPI (blue).

In all age groups, polymegathism was significantly increased in Q455K versus WT mice (*P<*0.0001), demonstrating a similar trend to ECD (**Figure 1C**). At 3 months of age, polymegathism began at 29.3±7.6% in the Q455K mice versus 16.7±1.4% in the WTs (*P*<0.0001). Within the WT group, the 24-month age group had significantly greater polymegathism than all other age groups (*P<*0.05). By contrast, Q455K mice demonstrated a significant increase in polymegathism between 3 months of age and all other ages (*P<*0.0001). Polymegathism was negatively correlated with ECD in both the WT and Q455K groups (*r*=−0.43, *P=*0.0008 and *r*=−0.84, *P*<0.0001 respectively; **Figure 1D**). Zonula occludens 1 (ZO-1) is a critical component of tight junctions and thus essential to the barrier function of CEnCs. In all mice ≤6 months of age, immunofluorescent ZO-1 staining was localized to the apical junctions of CEnC borders highlighting their normal hexagonal shape (**Figure 3C-D**). While WT mice ≥9 months of age maintained their hexagonal arrangement, the age-matched Q455K mice demonstrated cytoplasmic and apical junctional ZO-1 expression as well as progressively more severe polymegathism and pleomorphism consistent with what was observed with *in vivo* confocal microscopy (IVCM; **Figure 3**).

### Progressive guttae formation and quantification correlates with age and CEnC loss in Q455K mice

In the Q455K group only, guttae were identified as early as 3 months of age (29.2±22 guttae/mm^2^) and significantly increased in density to 273±37.3 guttae/mm^2^ by 24 months of age (*P<*0.0001, **Figure 4**). The guttae area ratio (GAR, expressed as percentage) increased significantly (*P<*0.0001) from 3- to 24-months of age at 0.8±0.54% to 7.2±1.5% respectively and a high correlation between guttae density (GD) and GAR was observed (*r=*0.88, *P=*0.007). Both GD and GAR positively correlated with age (*r*= 0.9, *P*<0.0001 and *r*=0.87, *P*<0.0001) and polymegathism (*r*=0.77, *P*<0.0001 and *r*=0.81, *P*<0.0001) as well as negatively correlated with ECD (*r*=−0.76, *P*<0.0001 and *r*= −0.87, *P*<0.0001; **Figure 4** and **Supplementary Figure 3)**. Further analysis of the guttae anatomical distribution demonstrated that that the majority of guttae were found at the vertices of CEnCs rather than in the center of cells (**Figure 5)** in both Q455K mice (74.3% vs. 25.7%) and human FECD patients (78.2% vs. groups (*P* < 0.001), with no significant difference in anatomical distribution found between the 2 species (*P=*1, **Figure 5**).

**Figure 4.**
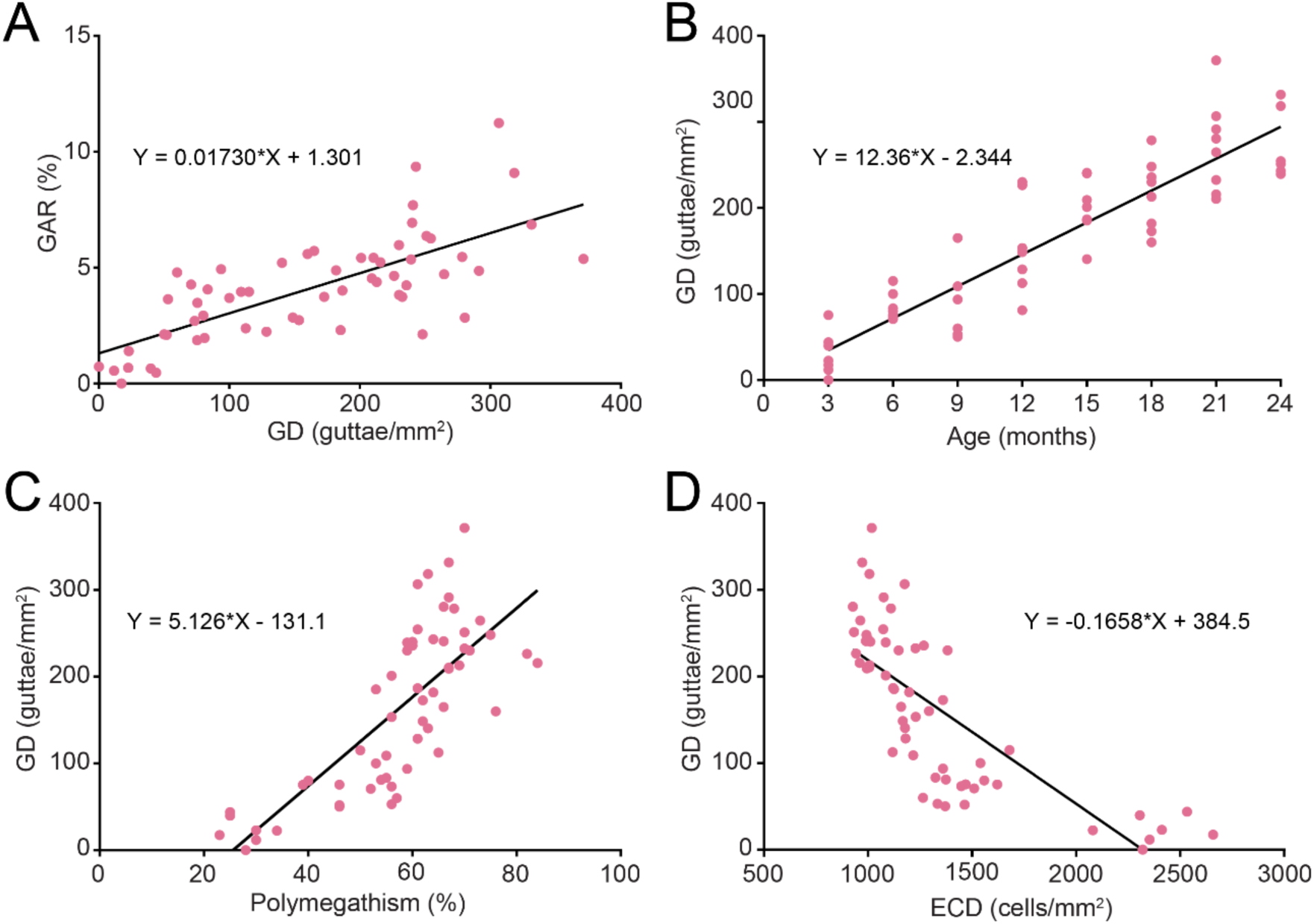
Guttae were exclusively observed in Q455K mice and increased with age. Given the high correlation between GD and GAR (*r=*0.88, *P=*0.007; **A**), GD only was used to determine correlations between age, ECD and polymegathism. The GD was positively correlated with age (*r*= 0.9, *P*<0.0001; **B**) and polymegathism (*r*=0.77, *P*<0.0001; **C**) as well as negatively correlated with ECD (*r*=−0.76, *P*<0.0001; **D**).

**Figure 5.**
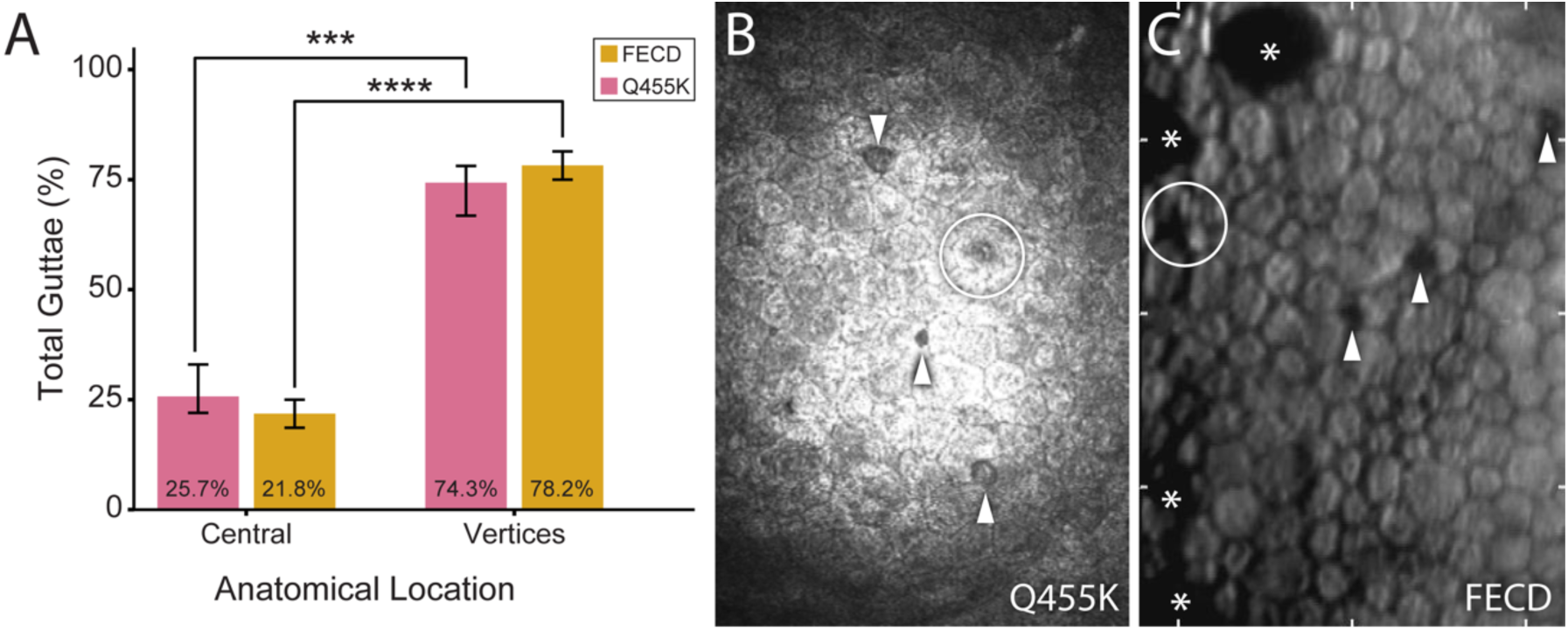
Guttae predominantly were identified at cell vertices in Q455K mice and FECD patients. Comparative analysis demonstrates (**A**) the regional percentage of total guttae localized to either the cell vertices or the central area in both human FECD patients (*P<*0.0001) and the Q455K murine model (*P<*0.001, initial guttae is concentrated at cellular junctions (78.2% and 74.3%, respectively) rather than centrally (21.8% and 25.7%). The spatial distribution pattern remains highly conserved and statistically indistinguishable between the 2 species (*P*> 0.05). In-vivo confocal microscopy image from a Q455K mouse (**B**) and a specular microscopy image from a FECD patient (**C**) demonstrating different guttae locations. Guttae location varies between the vertices (**arrowhead**) and the center (**circle**) of an endothelial cell. The asterisk (**C**) marks a more advanced guttae in an FECD patient that is either one large guttae or multiple that have coalesced. \*\*\**P<*0.001 and \*\*\*\**P<*0.0001.

### Upregulation of ferroptosis markers in CEnCs particularly surrounding guttae

Of the ten different genes associated with CEnC physiology (*Tjp1, Cdh2, Atp1a1, Atp1b2*) and ferroptosis (*Tfrc, Trf, Slc40a1, Ftl1, Fth1, Gpx4*) investigated, *Tfrc, Slc40a1, Ftl1, Gpx4,* had significantly different transcription levels between at least 2 groups, most commonly between WT and Q455K mice in the older age groups (**Figure 6**). Specifically, *Tfrc* and *Slc40a1* were significantly upregulated in Q455K mice 3-4 and >15 months of age versus WT mice >15 months of age (*P*≤0.04). Both *Tfrc* and *Slc40a1* encode trans-membrane proteins (transferrin receptor 1 or TFR1 and ferroportin respectively) responsible for transporting iron into (TFR1) and out (ferroportin) of the cell. Ferritin (FTL), encoded by *ftl1*, is the major protein responsible for iron storage within the cell and was significantly higher in Q455K mice >15 months of age in comparison both WT age groups (*P*≤0.026). In mice >15 months of age, *Gpx4* was significantly greater in Q455K versus WT mice (*P=*0.0284). GPX4, encoded by *Gpx4*, is the major enzyme responsible for protecting the cells against lipid peroxidation and thus, ferroptosis.

**Figure 6.**
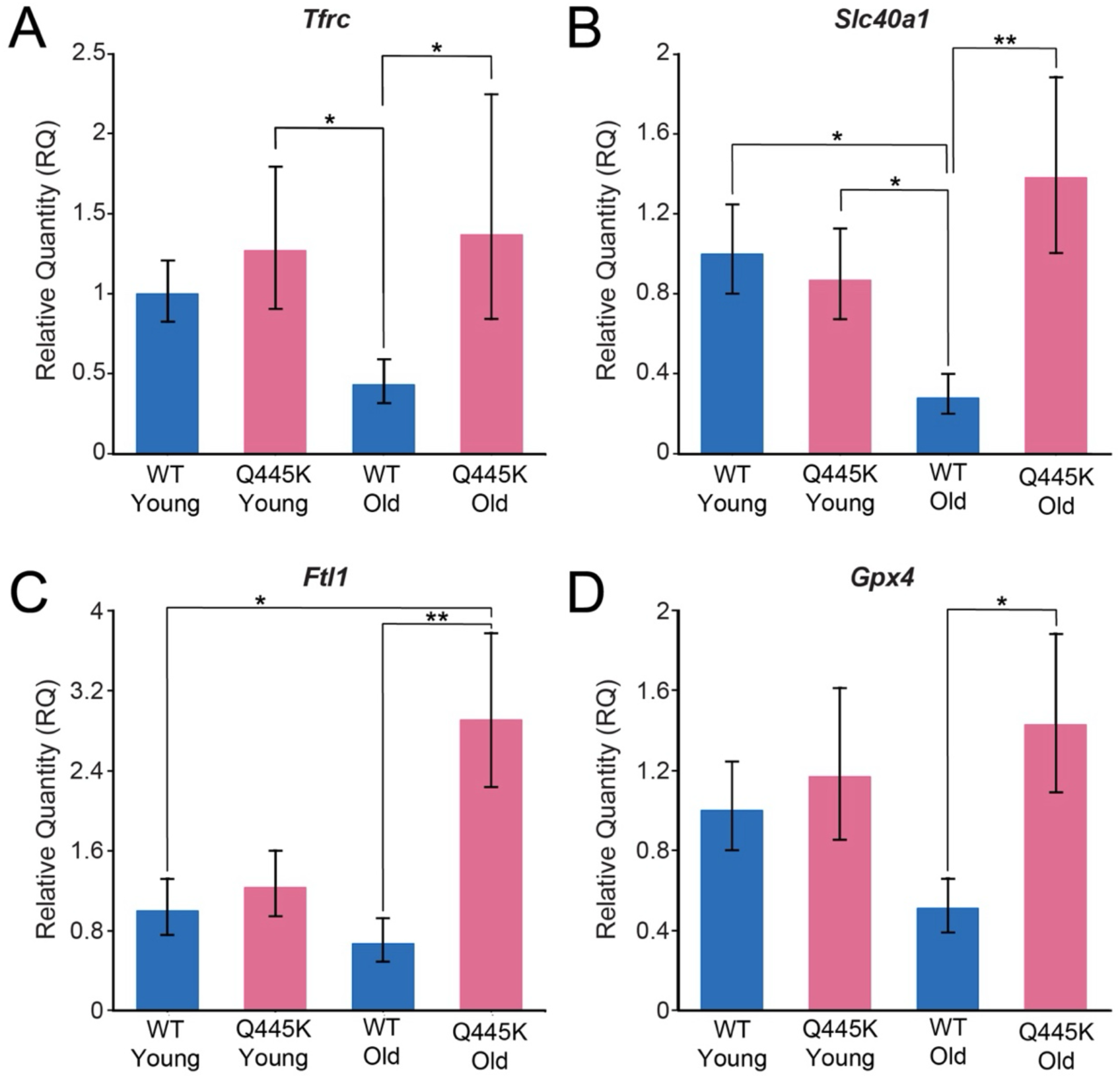
The mRNA expression of the ferroptosis-related genes, *Tfrc, Slc40a1, Ftl1,* and *Gpx4,* are upregulated in Q455K versus WT mice particularly at >15 months of age. The genotypes were divided into 2 age groups of 3-4 (young) or 15-25 (old) months. Expression of *Tfrc* was significantly greater in both the young and old Q455K mice versus the old WT controls (*P=* 0.04 and *P=*0.03 respectively; **A**). The *Slc40a1* expression was significantly greater in the young versus old WT mice (*P=*0.01), it was also significantly higher in the young and old Q455K mice when compared to the old WT controls (*P=*0.02 and *P=*0.001 respectively; **B**). *Ftl1* expression was significantly greater in the old Q455K mice versus young and old WT mice (*P=*0.03 and 0.002, respectively; **C**). Finally, *Gpx4* showed a significantly increased expression between old Q455K and WT mice (*P=*0.03; **D**). The ΔCt values were calculated between groups, normalizing genes of interest to 18S. P-values were calculated based on the ΔCt values. \**P<*0.05, \*\**P<*0.01, \*\*\**P<*0.001, and \*\*\*\**P<*0.0001, ns is not significant.

Protein expression for these 4 markers of ferroptosis also demonstrated upregulated expression in Q455K versus WT mice as well as in FECD versus non-FECD donor corneas with IHC (**Figures 7-9**). Specifically, ferroportin expression was more robust in Q455K murine and FECD human corneas and accumulated in the perinuclear space compared to normal controls (**Figure 7A-D**). The TFR1 expression was also upregulated in Q455K and FECD corneas compared to WT murine and non-FECD donor corneas, respectively (**Figure 7E-H**) with increased nuclear and cytoplasmic expression in CEnCs adjacent to guttae (**Figure 7F1-G2**). In addition, FTL demonstrated similar changes with increased cytoplasmic expression in Q455K versus WT mice (**Figure 8A-B**). By contrast, the non-FECD donor corneas demonstrated an increased cytosolic expression compared to the FECD donor corneas (**Figure 8C-D**). Lastly, GPX4 was more abundant in the cytoplasm of Q455K compared to WT mice (**Figure 9A-B**) as well as in FECD versus non-FECD donor corneas (**Figure 9C-D**) with greater nuclear expression in CEnCs adjacent to guttae (**Figure 9B1-C2**). To further confirm these distribution patterns, regional IHC quantification was performed on the Q455K mice. This analysis confirmed our localized observations, demonstrating that the mean intensity for all 4 ferroptosis markers (ferroportin, TFR1, FTL, and GPX4) were significantly elevated (*P<*0.05) in the CEnCs immediately surrounding guttae compared to the cells remaining in the image (**Figure 10**). In the human FECD patients, a similar statistical analysis could not be performed due to the high density of guttae (e.g. **Figure 7C1-C2**).

**Figure 7.**
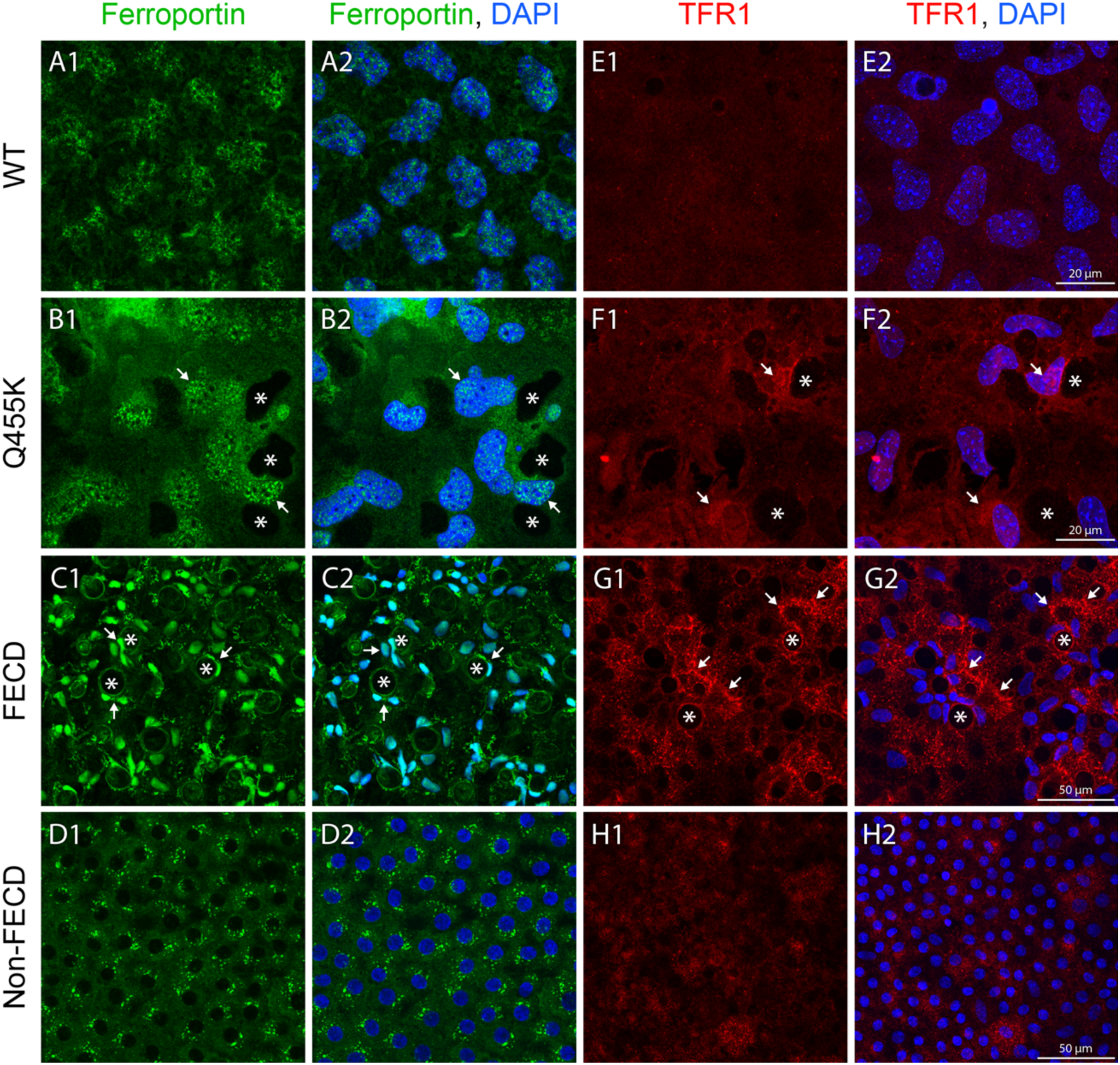
Perinuclear ferroportin (A-D) and transferrin receptor 1 (TFR1; E-H) were increased in the Q455K versus WT mice as well as in FECD corneas compared to non-FECD corneas. The Q455K mice and FECD patients also exhibited greater nuclear TFR1 and ferroportin in corneal endothelial cells (CEnCs) surrounding guttae (white arrows). Representative CEnC flatmounts from WT and Q455K mice at 21 months of age stained for ferroportin (green; **A1-B2**) and transferrin receptor 1 (TFR1, red; **E1-F2**), and nuclei (DAPI, blue). Similar CEnC flatmount preparations from FECD and non-FECD donor corneas were stained for ferroportin (**C1-D2**) and TFR1 (**G1-H2**). Increased staining intensity is highlighted with white arrows and representative guttae are marked with white asterisks. Scale bar represents 20μm for the murine corneas compared to 50μm for the human corneas.

**Figure 8.**
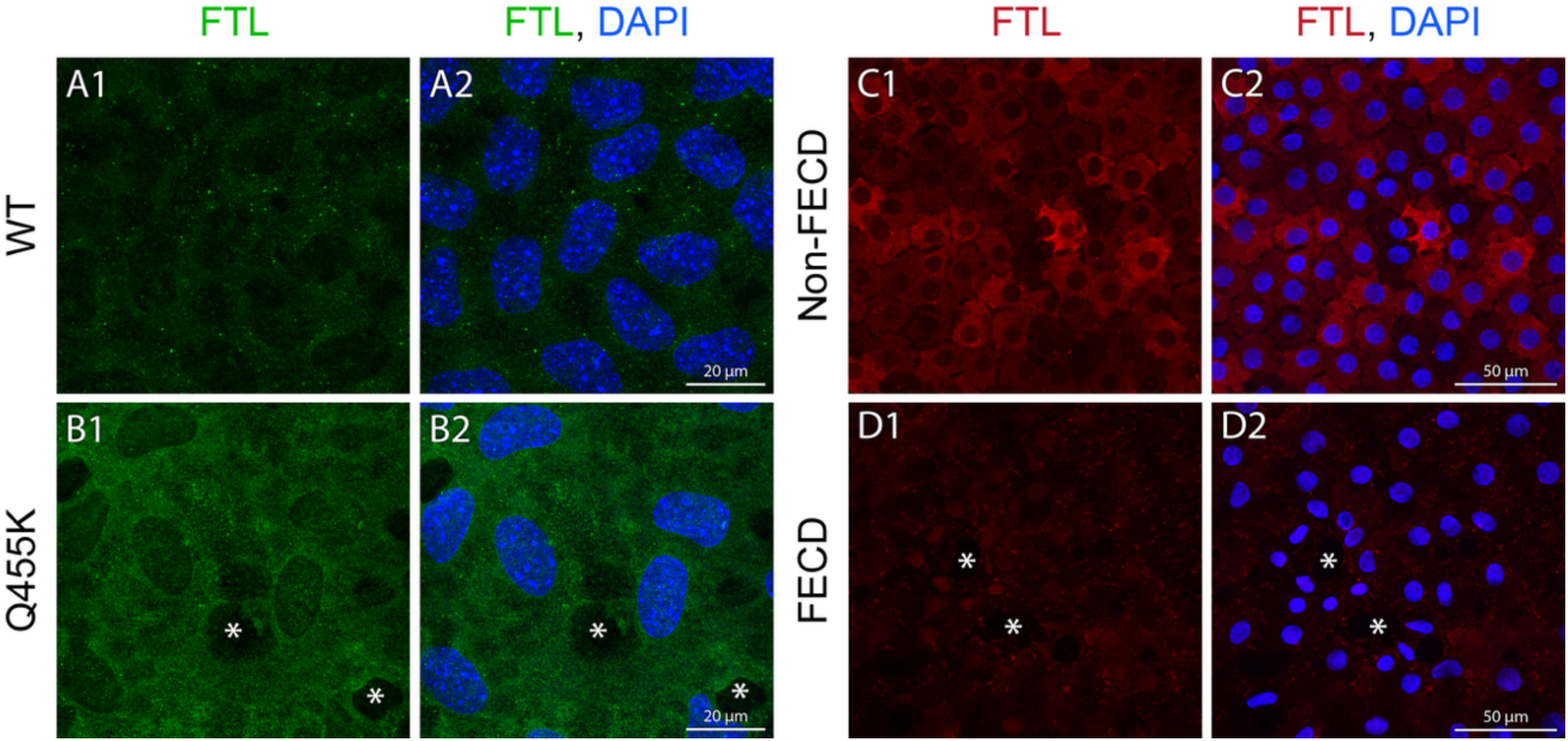
The WT mice exhibit less cytoplasmic ferritin (FTL) expression versus Q455K mice. By contrast, non-FECD corneas demonstrate higher FTL expression compared to FECD corneas. Representative CEnC flatmount preparations from WT and Q455K mice (**A1-B2**) at 21 months of age stained for FTL (green) and nuclei (DAPI, blue), as well as non-FECD and FECD donor corneas (**C1-D2**) stained for FTL (red) and nuclei (DAPI, blue). Guttae are marked with the white asterisks. Scale bar represents 20μm for the murine corneas versus 50μm for the human corneas.

**Figure 9.**
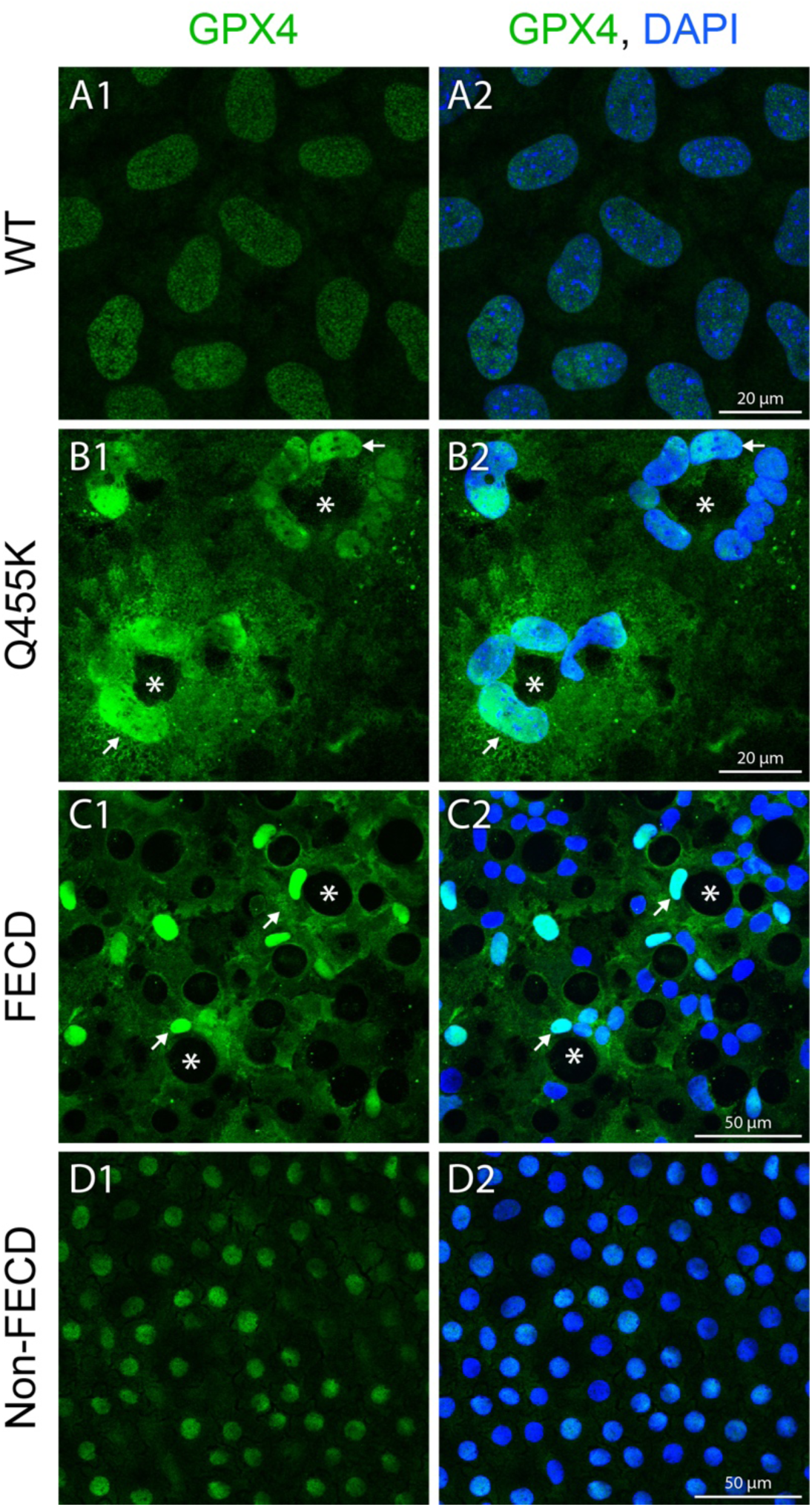
Glutathione peroxidase 4 (GPX4), is abundant in CEnC nuclei of WT mice while greater cytoplasmic and nuclear expression is observed in Q455K mice particularly adjacent to guttae (white arrows). Greater cytoplasmic and nuclear expression especially around guttae were also observed in FECD patients versus only nuclear expression in non-FECD controls. Representative CEnC flatmounts from Q455K and WT mice (**A1-B2**) at 21 months of age, as well as FECD and non-FECD donor corneas (**C1-D2**) staining for GPX4 (green) and nuclei (DAPI, blue). Guttae are marked with white asterisks and increased staining intensity is denoted with arrows. Scale bar equates to 20μm for the murine corneas compared to 50μm for the human corneas.

**Figure 10.**
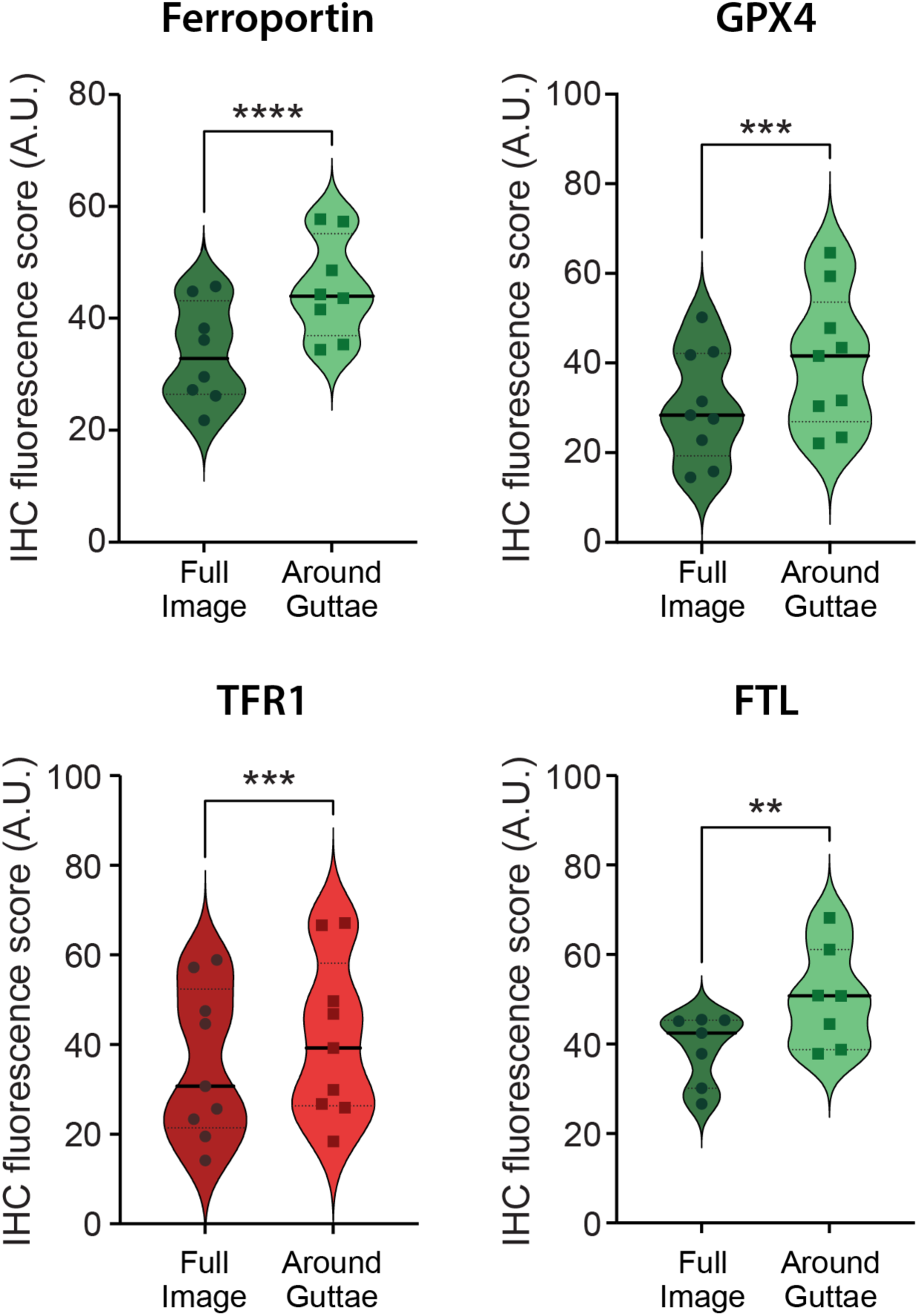
Quantification of fluorescence of ferroptosis markers was significantly higher in CEnCs surrounding guttae versus the remaining CEnCs in the Q455K mice (*P*<0.05). For each ferroptosis marker (ferroportin, GPX4, TFRC and ferritin) the intensity of the antibody florescence was measured specifically around the guttae region (light color) and for the entire image (dark color). \**P<*0.05, \*\**P<*0.01, \*\*\**P<*0.001, and \*\*\*\**P<*0.0001, ns is not significan

## Discussion

This study provides a comprehensive longitudinal evaluation of corneal disease progression and examines the role of ferroptosis in a murine model of early-onset FECD. By employing a combination of *in vivo* imaging and immunohistochemistry, we demonstrate that Q455K mice exhibit a progressive decline in ECD and progressive increases in polymegathism, guttae formation, and ferroptosis marker expression beginning as early as 3 months of age. These phenotypic changes are consistent with hallmark features of human FECD, provide critical evidence that ferroptosis is related to the presence of guttae and can occur across multiple genetic causes of FECD, and reinforce the translational utility of the Q455K murine model for preclinical therapeutic studies. Our findings confirm prior characterizations of the Q455K model (3, 4, 9), demonstrating not only the presence of FECD-associated features but also the timing and trajectory of disease progression over a 24-month period. As expected, we show that the decrease in ECD observed in Q455K mice is more rapid and severe than that observed with aging in WT controls. This supports previous observations that the Q455K mutation induces structural instability in the DM–endothelium interface, contributing to early CEnC reduction (4, 7).

Crosstalk between ECM remodeling and ferroptosis is a critical driver of disease progression in breast cancer, lung fibrosis, and liver fibrosis (24–26). This interaction is characterized by ECM proteins acting as regulators of the ferroptosis mechanism and, conversely, ferroptotic processes triggering excessive ECM deposition. In the present study, we observed upregulation of ferroptosis-related genes, *Tfrc*, *Slc40a1*, *Ftl1*, and *Gpx4*, and their proteins in in Q455K mice. These proteins are central regulators of iron metabolism and lipid peroxidation, key hallmarks of ferroptosis (20, 21). These data align with previous reports that human CEnCs in FECD are particularly susceptible to oxidative stress due to impaired antioxidant defense systems (18, 19). Reduced GPX4 expression and altered ferritin dynamics have been identified as contributing factors to ferroptotic susceptibility in FECD tissues (18). Furthermore, ferroportin, a key iron efflux protein, increases in human CEnCs with FECD (18). Similarly, the Q455K mice demonstrate increased ferroportin expression as well as localization to the perinuclear space, indicating excessive iron transport thus supporting the presence of ferroptosis. Although ferritin was decreased in CEnCs with late-stage FECD (18) it was increased in the Q455K mice in the present study. This increase would be expected early in disease in order to chelate ferrous iron to prevent Fenton chemistry, lipid peroxidation of the plasma membrane, and ultimately, ferroptosis (27, 28). A major player in preventing lipid peroxidation that leads to ferroptosis is GPX4 and both its mRNA and protein expression is increased in the Q455K mice. Furthermore, more GPX4 is localized in the nuclei of CEnCs adjacent to guttae. The localization pattern differences of ferroptosis markers between WT versus Q455K mice as well as non-FECD versus FECD corneal tissues represent a potential isoform prevalence shift in response to increased FECD pathology and a topic for future study (22). Notably, our findings reveal that the fluorescence intensity of ferroptosis markers is significantly higher immediately adjacent to guttae compared to CEnCs further away. This observation highlights the intense metabolic and oxidative stress concentrated at the borders of these DM excrescences and may act as key initiators of ferroptosis. Given the growing recognition of ferroptosis in CEnC pathophysiology, these results further support the study of ferroptosis inhibitors such as lipophilic antioxidants or iron chelators as potential therapeutic strategies to slow or prevent disease progression in FECD (23, 29, 30).

Initial guttae location appears strikingly similar between Q455K mice and humans with FECD with ∼75-80% at cell vertices in both groups. Within the Q455K and FECD tissues, CEnCs adjacent to guttae demonstrate a similar pattern of increased intensity of ferroportin, TFR1, FTL and GPX4. Knockdown of GPX4 in human CEnCs increases lipid peroxidation, cytotoxicity, and susceptibility to iron- and peroxide-induced injury (31). The elevated GPX4 expression we detected in Q455K murine and FECD patient corneas likely represents an initial attempt to mitigate the ferroptotic pathway and preserve endothelial viability. It is notable that all four ferroptosis markers investigated demonstrate upregulated expression surrounding guttae suggesting that guttae may initiate iron dysregulation and endothelial stress.

Consistent with our observations, Saha et al. reported that FECD corneal tissues exhibiting confluent guttae display elevated Fe²⁺ accumulation and lipid peroxidation, indicative of increased ferroptosis surrounding guttae (18). These findings parallel those of Méthot and colleagues who demonstrated that in human FECD explants, CEnCs adjacent to guttae exhibit reduced mitochondrial potential and mass, increased intra-mitochondrial calcium, and greater cell loss, indicating that guttae impose metabolic and oxidative stress on neighboring cells (13). Our data extend these findings by implicating ferroptosis as a mechanistic link between guttae-associated mitochondrial dysfunction and CEnC death, now in a second and distinct FECD disease-causing genetic mutation. Together, these findings highlight ferroptosis as a cellular death pathway contributing to localized endothelial degeneration around guttae, reinforcing their role as focal microenvironments of oxidative injury in FECD.

The presence and quantification of guttae formation in this study further strengthen the relevance of the Q455K mouse as a model of FECD. In TAZ (*Wwtr1*) deficient mice, a late-onset FECD model, guttae are absent although this model demonstrates DM softening suggesting that ECM pathology is still present (32). Unsurprisingly, guttae are present in a new double mutant mouse bearing both the *Col8a2* Q455K knock-in and a tamoxifen-inducible Slc4a11 knockdown (11). Herein, we demonstrate that guttae density and GAR increased significantly with age and highly correlate with polymegathism and ECD. This is notable as guttae are considered principally indicative in FECD and thought to originate from aberrant ECM secretion and cellular stress responses (1, 33–35). In humans, disease severity is primarily linked to guttae size, as typically assessed via the Krachmer scale during slit-lamp examination (6, 14, 36, 37). Due to the marked size difference between human and murine corneas, this scale is less applicable in mice, prompting alternative measures of guttae severity. Although GAR has been shown to correlate with modified Krachmer grading (16, 34), we observed that guttae increased predominantly in their quantity rather than size (e.g GAR), suggesting guttae density may better reflect disease severity in this murine model (38). Thus, guttae density can be used as an objective parameter in murine models to assess progression and treatment response. Additionally, literature regarding the morphological classification of guttae in FECD remains limited. In a foundational study, Son and colleagues categorized guttae into 5 distinct stages characterized by size, cellular abnormalities, coalescence, and contour, noting that multiple stages frequently coexist within a single cornea (39, 40). Expanding upon structural characterizations, our analysis of anatomical distribution revealed a distinct spatial/topographical pattern, demonstrating that the majority of total guttae were found at the cell vertices rather than the cell center both in the Q455K mouse model and FECD patients.

Interestingly, despite the marked reduction in ECD, none of the Q455K mice developed corneal edema as evidenced by increased CCT by 24 months of age. This finding suggests that corneal endothelial pump function is preserved to a critical threshold likely via upregulation of Na⁺/K⁺-ATPase combined with the rodents’ endothelial proliferative capacity (41). Furthermore, observations in human and canine patients suggest that corneal edema typically manifests only after ECD drops below ∼500 cells/mm² (41–43) and all mice maintained a cell density >900 cells/mm² until 24 months of age in the present study. While the absence of edema reflects a common limitation in murine FECD models, this occurs at the end-stage of FECD following the optimal period by which medical therapeutic interventions would be optimally administered (44). It is also possible that corneal edema could be generated in this model with ultraviolet-A induced damage to accelerate CEnC loss (18, 45, 46).

Polymegathism, a measure of CEnC stress along with pleomorphism, was significantly increased in Q455K relative to WT mice at all timepoints and demonstrated a strong negative correlation with ECD. This is consistent with findings in human FECD and further supports the fact that polymegathism precedes endothelial decompensation (1, 5, 7, 37, 41). Additionally, immunostaining revealed cytoplasmic expression of ZO-1, suggesting compromised barrier integrity. This loss of ZO-1 hexagonality is consistent with endothelial stress and has been documented in human FECD specimens as well as other murine models of FECD (1, 3, 4, 7, 10, 11, 32).Together, these findings highlight ZO-1 disruption as an objective, quantifiable indicator of polymegathism and pleomorphism (i.e. early dysfunction), providing valuable metrics for evaluating disease progression and the efficacy of targeted therapeutics in this model.

The primary limitation of this study was the inability to follow an individual mouse throughout the full 24-month period to track disease progression. Due to natural lifespan constraints and housing limitations, different cohorts of mice were used at various time points.

In conclusion, the *Col8a2* Q455K mouse recapitulates key structural and functional aspects of early-onset and late-onset FECD, including early and progressive ECD loss, polymegathism, and guttae formation. This study provides robust longitudinal data on disease progression, supporting the use of this model in evaluating targeted therapeutics, particularly those aimed at preserving CEnC function and/or preventing ferroptosis. This model can be utilized to dissect the molecular pathways involved in FECD, particularly ferroptosis, to further elucidate mechanisms of CEnC loss and intervention points for medical therapy to prevent FECD disease progression.

## Materials and Methods

### Animals

All studies were approved by the Institutional Animal Care and Use Committee of the University of California-Davis and performed according to the Association for Research in Vision and Ophthalmology resolution on the use of animals in research. The Q455K and WT mice were bred and genotyped as previously described (7), and evaluated from 3 months of age, every 3 months, for a total period of 24 months. Each age group had between 6-8 mice. For IHC staining of ZO-1 and Na^+^/K^+^-ATPase, we included 6 mice from 4 age groups (3,6, 12 and >18 months). For ferroptosis analysis, we included mice from either genotype (WT or Q455K) for 2 age groups (3-4 and 15-24 months) for qPCR (n=8-11/group) and IHC analysis (n=10/group).

### Human Tissue Consent and Collection

All investigations were carried out following the guidelines of the Declaration of Helsinki. Tissues were obtained with informed consent by the donor’s family or next of kin for donor control tissues or by the patient undergoing keratoplasty for FECD tissues. Approval was determined as not required for the deidentified donor corneal tissues according to the Institutional Review Board (IRB) at the University of Iowa. For FECD samples, human corneal endothelial tissue was collected at the University of Iowa at the time of endothelial keratoplasty from patients with advanced FECD that were enrolled in the Proteomic Analysis of Corneal Health Study (PACH IRB #201603746). For control samples, human corneal endothelial tissue was obtained from human donor eyes provided by the Iowa Lions Eye Bank (ILEB, Coralville, IA) and Beauty of Sight (Miami, FL).

### Sex as a Biological Variable

Both male and female Q455K mice were included in this study, and potential sex-specific differences were evaluated for ECD, polymegathism, GD, and GAR. Human clinical samples were also obtained from both male and female subjects. Due to a baseline asymmetry in sample distribution between the cohorts (predominantly female in the diseased group versus predominantly male in the control group), comparative statistical analysis between sexes was not performed.

### Ophthalmic examinations and multimodal corneal imaging

Mice were manually restrained for an ophthalmic examination performed with a handheld slit lamp (SL-17, Kowa, USA). Clinical findings were recorded and scored using the Semiquantitative Preclinical Ocular Toxicology Scoring (SPOTS) system (47). For imaging, mice were sedated with an intraperitoneal injection of ketamine (75 mg/kg) and dexmedetomidine (0.5 mg/kg). Slit lamp biomicroscopy with digital capture (Hagg-Streit BQ 900 Slit Lamp; Hagg-Streit) was performed to photograph the eyes. In vivo confocal microscopy (IVCM) was performed with the Heidelberg Retina Tomograph III Rostock Corneal Module (HRT3, Heidelberg Engineering GmbH, Heidelberg, Germany) confocal microscope was utilized to assess corneal ECD and morphology, with 3 representative regions counted for each eye. Using ImageJ, a region of interest (ROI) was chosen (>0.03) and cells were marked and counted to calculate ECD, mean cell area and polymegathism. Within the Q455K group, we calculated guttae density by dividing the number of guttae by the ROI. Guttae area ratio (GAR) was calculated as previously described using the following equation (34, 38):

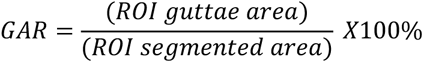

Fourier-domain optical coherence tomography (FD-OCT; RTVue 100, software version 6.1; Optovue, Inc., Fremont, CA, USA) with a CAM-S lens (3-mm scan length) was utilized to measure CCT via imageJ with 4 representative regions counted for each eye (2 horizontal and 2 vertical images). The center of the scan pattern was aligned with the corneal vertex reflection (48) visualized on the FD-OCT images (**Figure 2**).

The ocular surface was kept moist using a lubricant (GenTeal^Ⓡ^ Tears; Alcon; Fort Worth, Texas) during all image modalities. Guttae location analysis was performed on IVCM images from the 3-month-old mice and specular microscopy images from FECD patients with a Krachmer grade of ≤5.

### Ferroptosis qPCR analysis

Mouse corneas were carefully dissected along the limbus and placed endothelium side up as described by Uehara et al.(49). Care was taken to avoid inclusion of the iris, trabecular meshwork, and ciliary body. Using fine non-toothed microforceps under a stereomicroscope, DM was gently grasped at the peripheral edge and peeled centripetally toward the central cornea. Corneas collected from both eyes of each mouse were pooled and placed into 200 μL of extraction buffer from the PicoPure RNA Isolation Kit (PicoPure RNA isolation kit, Cat# KIT0204; Thermo Fisher Scientific, Carlsbad, CA,USA) then immediately snap-frozen and stored at −80 °C until RNA extraction.

RNA was purified with the PicoPure RNA isolation kit following the manufacturer’s guidelines and quantified using the Qubit 4 fluorometer using Qubit RNA high sensitivity kit (Cat# Q32852; Thermo Fisher Scientific,). Total RNA (20 ng) was reverse transcribed in 40 µL reaction volume using a High-Capacity cDNA Reverse Transcription kit (Cat# 4368814, Applied Biosystems/Thermo Fisher Scientific). Total reaction volume in qPCR assay was 20 µL containing 10 µL 2x SsoAdvanced Universal SYBR Green Supermix, 1 µL each of 10 uM forward and reverse primers (**Supplementary Table 1**), 7 µL nuclease-free water, and 1 µL of cDNA template. Real-time qPCR was performed on the CFX Connect thermal cycler (Bio-Rad Laboratories) with the following cycling parameters: 30 seconds at 95°C for initial denaturation, then 40 cycles of 10-second denaturation step at 95°C and 30-second annealing/extension step at 60°C. Immediately after amplification, a melt curve protocol was performed between 65°C and 95°C with 5 seconds each at 0.5°C increments. Melting curves were generated and single product formation was confirmed for all primer pairs used and all assays. The ΔCt values were calculated between groups, normalizing genes of interest to 18S. The ΔΔCt values were calculated between groups of interest and then subsequently used to calculate relative quantity (RQ; 2^−(ΔΔCt)^) and their confidence intervals bounded by RQ_min_ and RQ_max_.

### Immunohistochemistry (IHC)

To further assess CEnC structure and function, IHC was performed on WT and Q455K corneal flatmounts for key the CEnC markers, ZO-1 and Na^+^/K^+^-ATPase. Enucleated globes were collected for subsequent corneal dissections. Corneoscleral buttons were dissected and fixed in 4% paraformaldehyde (PFA; Santa Cruz Biotechnology, California) for 5 minutes with the endothelial side of the corneas facing upwards. After rinsing with phosphate buffered solution (PBS), they were immediately immersed into 0.1% Triton X-100 (SigmaAldrich; St. Louis, Missouri) for 1 hour at room temperature. Corneas were then placed into blocking solutions comprising 2% Bovine Serum Albumin (BSA; SigmaAldrich; St. Louis, Missouri), 5% normal donkey serum, 0.1% TritonX-100 all diluted in PBS for another hour at room temperature. Primary antibodies for the following targets were used: ZO-1 rabbit polyclonal antibody (Invitrogen; Carlsbad, California; 61-7300; 1:100) and Na^+^/K^+^-ATPase rabbit antibody (Invitrogen; MA5-32184; 1:200). Following overnight incubation at 4℃, corneas were then briefly washed with primary antibody diluent and 0.1% Triton-X for 15 minutes each. Corneas were then incubated with secondary antibodies AlexaFluor594 (LifeTechnologies; Carlsbad, California; A21207; 1:200) and AlexaFluor488 (Invitrogen; A21260; 1:200) for 1 hour at room temperature in the dark. Post-secondary antibody incubation, corneas were washed with PBS twice for 10 minutes each and then briefly stained with Hoescht 33342, trihydrochloride trihydrate (LifeTechnologies; Carlsbad, California; H3570) for nuclear staining. Following nuclear staining, corneas were flattened, cut into 4-6 leaflets and mounted down onto slides using ProLong Gold antifade reagent (Invitrogen; P36930).

Immunohistochemistry for proteins associated with ferroptosis was performed on mouse corneas and human surgical explant or donor tissues. Fixed mouse corneas were divided in half, and 3 radial cuts were made in each half-cornea for flat mounting later. Samples were incubated in 0.1% Triton X100/PBS solution for 30 min and then in 2% BSA/5% goat serum/0.1% Triton X100/PBS solution for 60 minutes to achieve permeabilization and blocking. Each sample was incubated in combination or with one of the following primary antibodies for 20 hours at +4°C and 0.1% BSA/0.5% goat serum/0.1% Triton X100/PBS solution used as primary antibody diluent: 1) rabbit monoclonal antibody against GPX4 (diluted to 10.4 µg/mL, Abcam #AB125066, Waltham, MA, USA) and rat polyclonal antibody against transferrin receptor 1 (TFR1, diluted to 10 µg/mL, BiCell Scientific #31013, Maryland Heights, MO, USA); or 2) rabbit polyclonal antibody against ferritin light chain (FTL, diluted to 20 µg/mL, Proteintech #10727-1-AP, Rosemont, IL, USA); or 3) rabbit polyclonal antibody against ferroportin (diluted to 33 µg/mL, Novus Biologicals #NBP1-21502, Centennial, CO, USA). Primary antibody solution was removed next day with 4 washes in 0.1% Triton X100/PBS. Alexa Fluor® 488 goat anti-rabbit Superclonal™ secondary antibody (diluted to 2.5 µg/mL, Invitrogen #A27034) and Alexa Fluor® 568 goat anti-rat secondary antibody (diluted to 2 µg/mL, Invitrogen #A11077) in 0.1% Triton X100/PBS supplemented with 300 nM DAPI (Invitrogen #D1306, Thermo Fisher Scientific, Waltham, MA, USA) was applied for 2 hours in darkness at room temperature.

Human tissue was placed in OptisolGS storage media (Bausch & Lomb, Irvine, CA) at 4°C immediately after collection and then fixed with 4% paraformaldehyde solution buffered with 0.1M PBS, pH 7.4 for 10 minutes at room temperature within 2-4 hours after surgery. Donor tissues without history of FECD were prepared by mounting donor corneas onto a 9.5 mm vacuum trephine (Barron Precision Instruments, LLC, MI, USA) and scoring the endothelium and Descemet membrane into the stroma. The endothelial Descemet’s membrane complex was visualized with 0.06 % trypan blue ophthalmic solution (VisionBlue, DORC International, Netherlands) and the tissue was carefully peeled away from the stroma and fixed with 4% paraformaldehyde solution buffered with 0.1M PBS, pH 7.4 for 10 minutes at room temperature within 2 weeks of tissue procurement. Fixed tissue was rinsed with 0.02M PBS, pH 7.4 up to 3 times and stored at 4°C in PBS until immunolabeling procedure. Permeabilization, blocking, primary and secondary antibody incubation conditions were the same as described for IHC on mouse cornea, except that a higher blocking agent concentration (0.2% BSA/1% normal goat serum) was used for overnight incubation with anti-TFR1 antibody. One of the following primary antibodies was applied to human tissue: 1) rabbit monoclonal antibody against GPX4 (diluted to 12.5 µg/mL, Abcam #AB125066, Waltham, MA, USA); 2) rabbit polyclonal antibody against ferritin (diluted to 10 µg/mL, Invitrogen #PA1-29381); or 3) rabbit polyclonal antibody against ferroportin (diluted to 33 µg/mL, Novus Biologicals #NBP1-21502, Centennial, CO, USA); 4) mouse monoclonal antibody against TFR1 (diluted to 2.5 µg/mL, Millipore-Sigma #MABS1982, Billerica, MA, USA). Ferroportin and GPX4 were detected with Alexa Fluor® 488 goat anti-rabbit Superclonal™ secondary antibody (2.5 µg/mL, Invitrogen #A27034), ferritin with AlexaFluor594 AffiniPure Fab fragment goat anti-rabbit antibody (16 µg/mL, Jackson ImmunoResearch #111-587-003, West Grove, PA, USA), and TFR1 with AlexaFluor568 goat anti-mouse antibody (2.0 µg/mL, Invitrogen #A11004). At the end of incubation with secondary antibody and DAPI, each sample was washed several times with 20 mM PBS, pH 7.4 and distilled water and mounted under Epredia™ Aquamount mounting medium (Epredia #13800, Thermo Fisher Scientific).

### Statistical analysis

Group comparisons were conducted by a 2-way ANOVA combined with Tukey’s multiple comparisons test, with individual variances for each comparison using GraphPad Prism (La Jolla, CA, version 10.4.2). Normality was determined for each data set by the Shapiro Wilk test for normality.

Bivariate associations were computed with Spearman rank correlation for nonparametric data and with Pearson correlation when normality was achieved. The *P* values for qPCR of ferroptosis markers were calculated based on the ΔCt values using Tukey’s multiple comparisons test in Graphpad Prism. *P* values less than 0.05 were considered statistically significant data are presented as mean ± standard deviation (SD). To determine if the topographic distribution of guttae observed in the murine model was representative of human pathology, a Chi-square test of independence was performed using R software (version 4.5.2; 10.31.2025, R Foundation for Statistical Computing, Vienna, Austria). The difference in IHC intensity between regions was assessed with a paired *t*-test.

## Data, Materials, and Software Availability

Anonymized data underlying all analyses and supporting the findings presented in this manuscript are available in a single Excel workbook (Col8_data_Anonymized.xlsx) deposited in the Figshare repository and are publicly accessible at (https://doi.org/10.6084/m9.figshare.32811098). The workbook contains the measurements used for all main figures and tables, including Q455K and WT ECD, polymegathism, CCT and guttae analysis, as well as IHC information for both human and mouse samples and human specular microscopy data used for guttae location analysis. No custom software was created for this study; statistical analyses were performed using GraphPad Prism (version 10.4.2) and R software (version 4.5.2) as described in the *Materials and Methods*.

## Acknowledgments

This research was supported by grants from the National Institutes of Health (NIH R01EY016134, R01EY037135, R01EY036440, R01EY033733-1A1S1, P30EY012576, T32GM13574). The authors have no conflicts of interest or disclosures related to this work. The authors thank the many members of the Comparative Ophthalmology and Vision Sciences Laboratory for their contributions throughout the study. We appreciate Ana Ripolles-Garcia for her guidance with statistical analysis. We are grateful to Chrisoula A. Toupadakis Skouritakis, Director of MediaLab Services at the School of Veterinary Medicine, for meticulous figure editing. The authors also wish to acknowledge the support of the Beulah and Florence Usher Chair in Cornea/External Disease and Refractive Surgery, Cornea Transplant Research Fund, Iowa Lions Eye Bank, M.D. Wagoner & M.A. Greiner Cornea Excellence Fund, and Seidler Foundation. We are grateful for the support of our patients, cornea donors, and donor families.

## Author Contributions

K.W.H. performed data collection, analyzed data, and wrote the initial draft of the manuscript. J.L. contributed to data collection and writing an initial draft paragraph. H.I. contributed to data collection and data analysis J.M.S. also assisted in writing and data curation. S.K., H.S., S.P., N.E., M.F., J.S and K.P.R. contributed to data collection. M.J.K., G.L.D., M.I., L.J.Y., M.A., J.M.S, M.A.G., and S.L. contributed to data analysis. B.C.L. contributed to study design and reviewed the final manuscript. M.A.G. reviewed the final manuscript. S.M.T. conceived the study, acquired funding, supervised the project and manuscript preparation. All authors edited, reviewed and approved the final manuscript.

## Supplementary Data

**Supplementary Figure 1.**
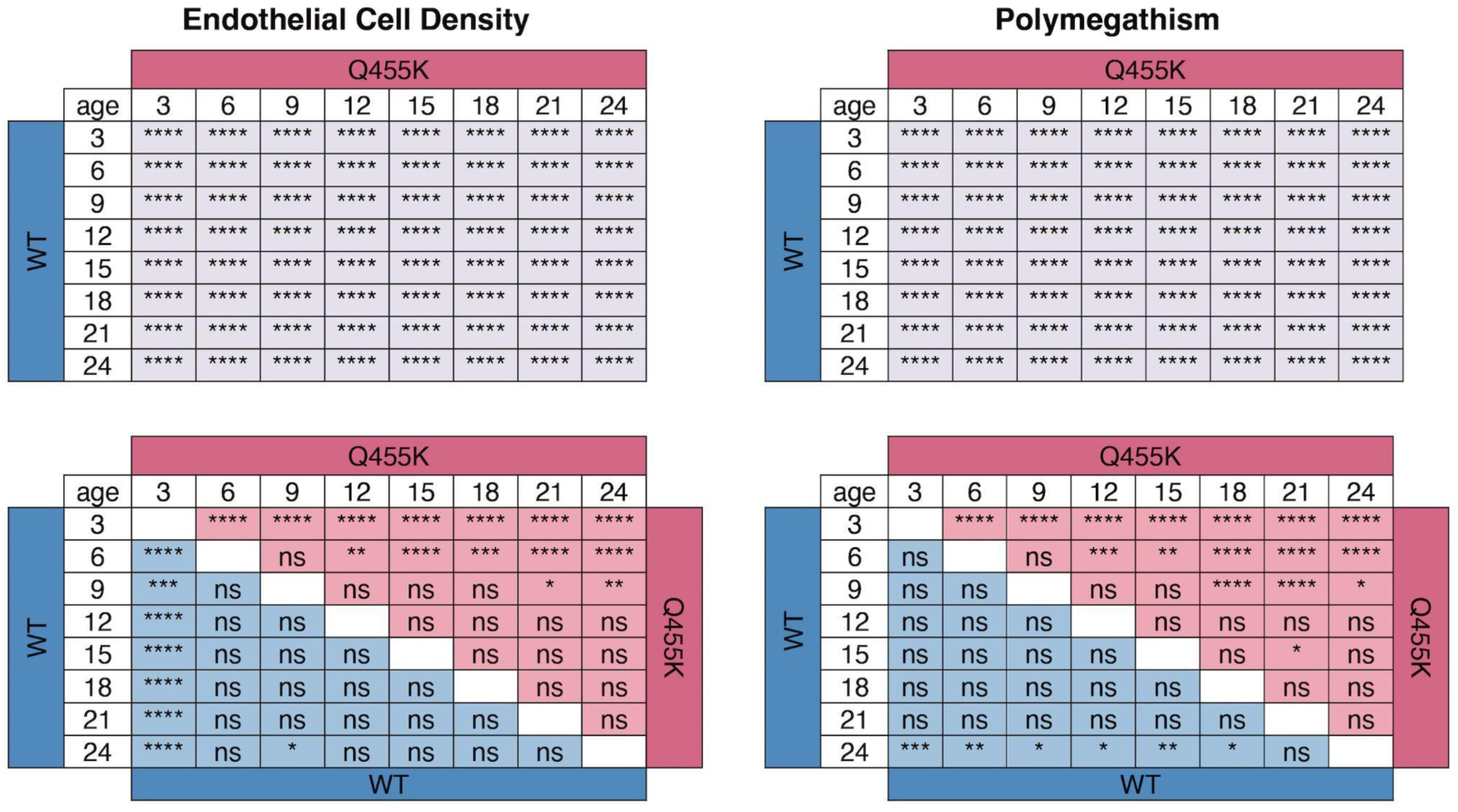
Matrix graphs denote *P* values of the 2-way ANOVA analysis for CEnC density and polymegathism. Tukey’s multiple comparisons test, with individual variances, was done for each comparison. \**P<*0.05, \*\**P<*0.01, \*\*\**P<*0.001, and \*\*\*\**P<*0.0001, ns is not significant.

**Supplementary Figure 2.**
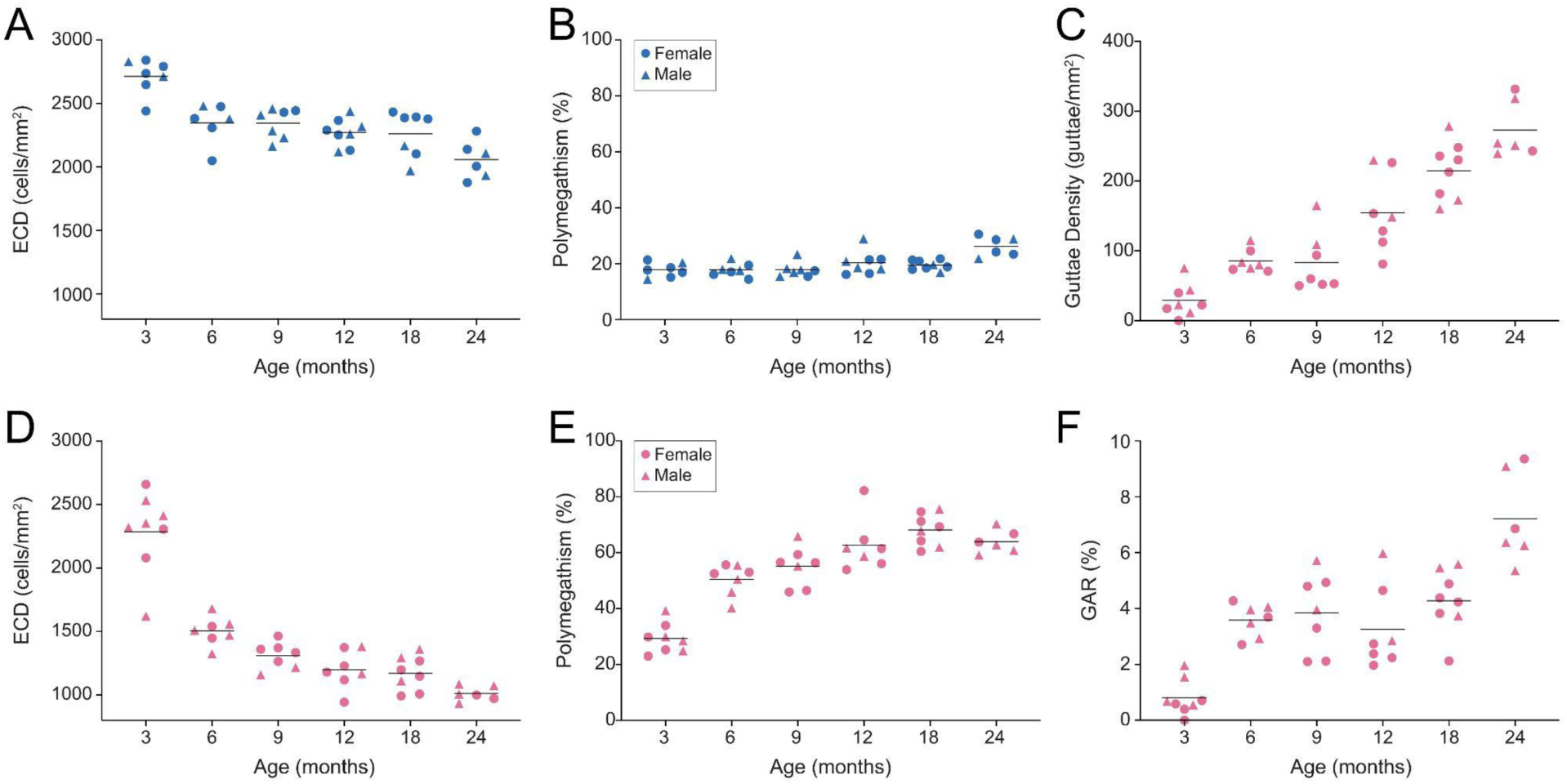
Endothelial cell density (ECD) and polymegathism did not significantly differ between sexes in WT (blue) or Q455K (pink) mice, guttae density (GD) and GAR% did not differ between sexes within the Q455K mice. The bars represent the mean value. The 15- and 21-month age groups were excluded due to a homogenous sex group.

**Supplementary Figure 3.**
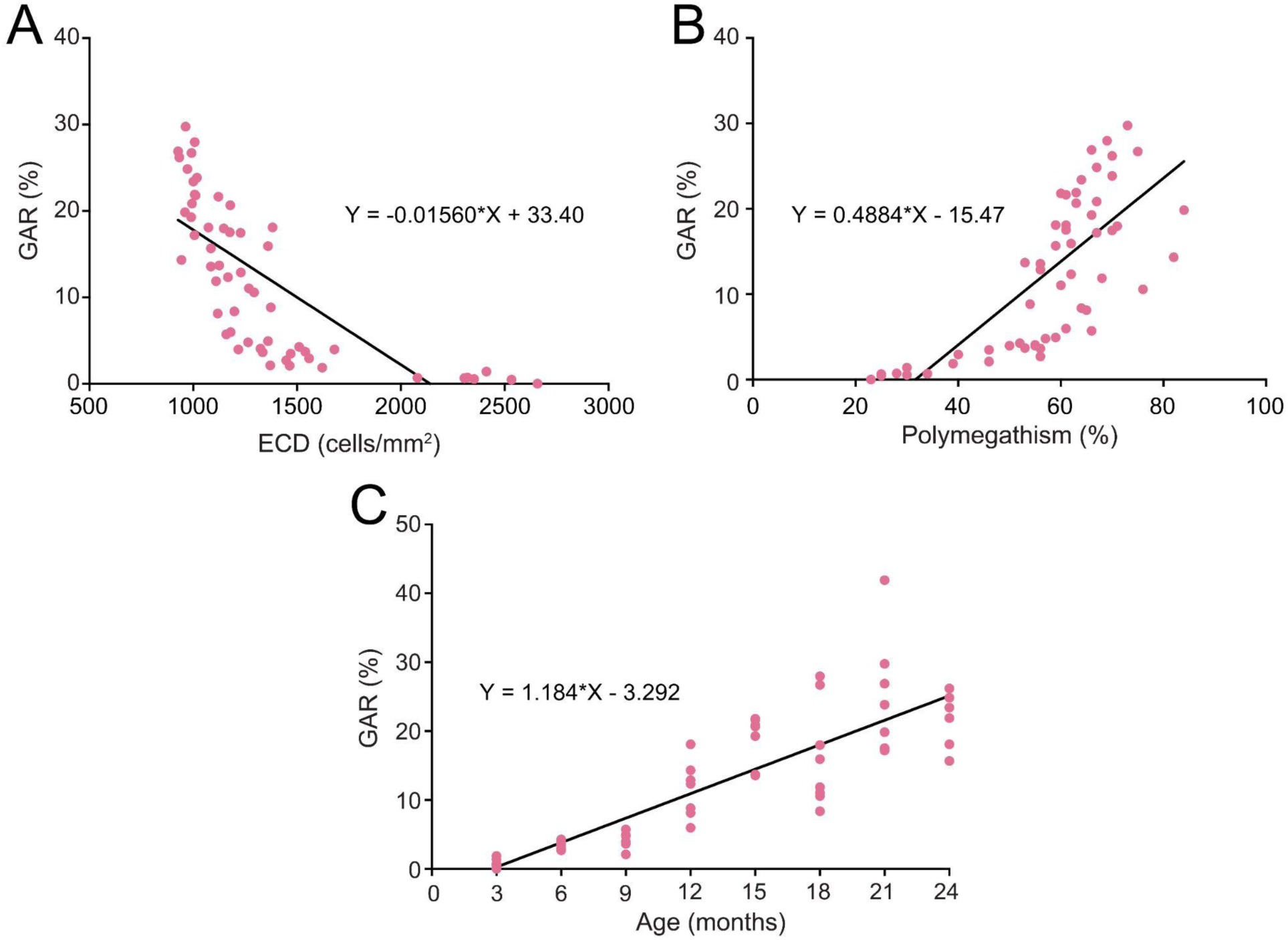
Guttae area ratio (GAR) was exclusively calculated in Q455K mice demonstrating the guttae phenotype. The GAR was negatively correlated with ECD (*r*= −0.87, *P*<0.0001; **A**) and negatively correlated with polymegathism (*r*=0.81, *P*<0.0001; **B**) and age (*r*=0.87, *P*<0.0001; **C**).

**Supplementary Table 1.**
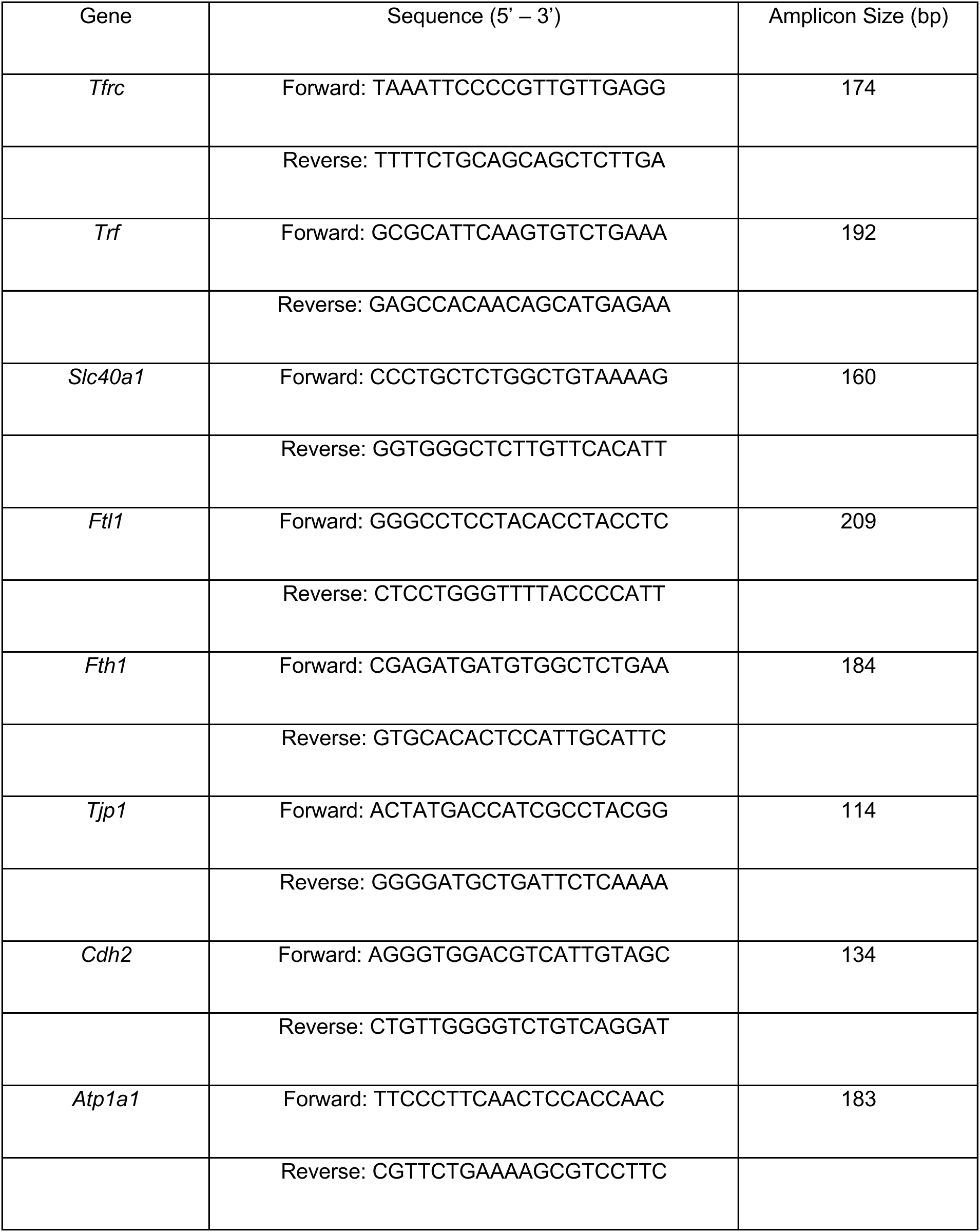
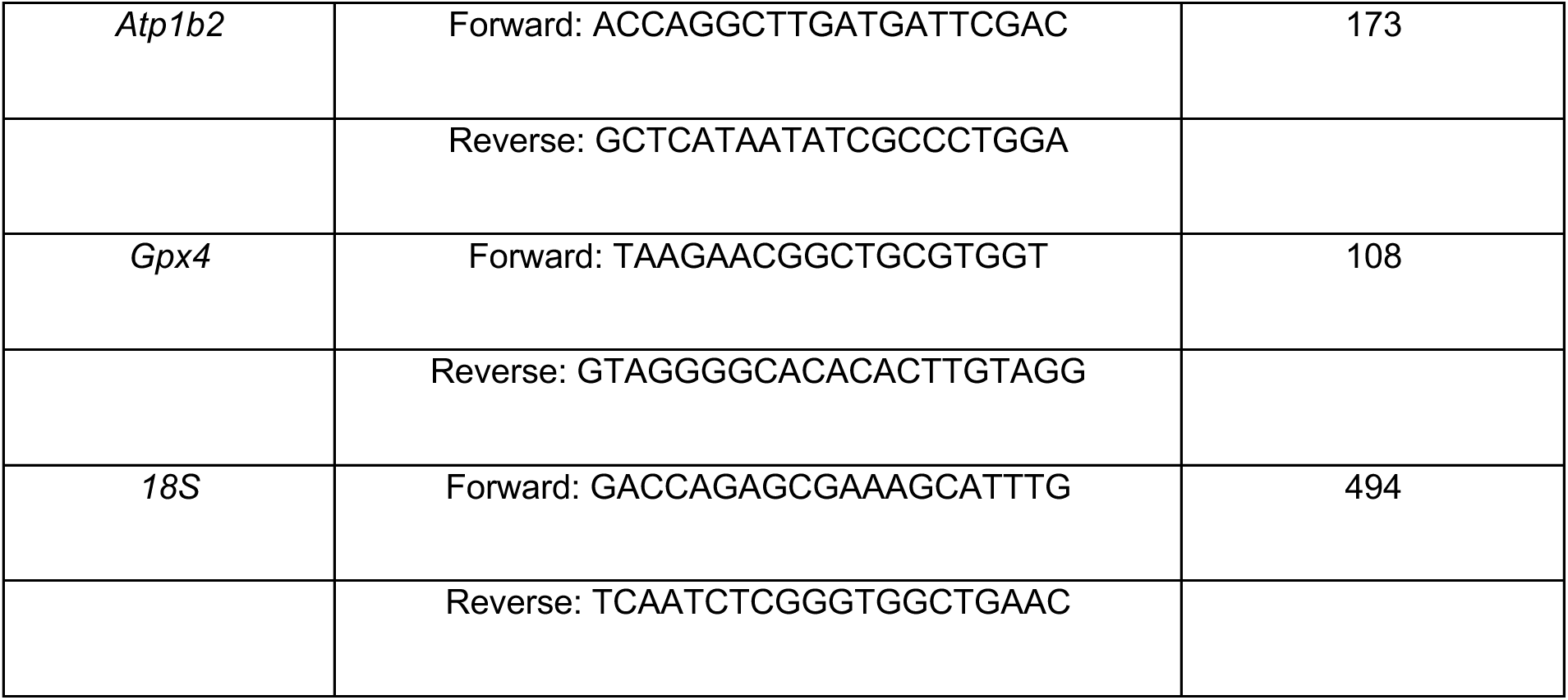
Primers used for mouse qPCR assays.

**Supplementary Table 2.** Individual measurements for Q455K mice including age, endothelial cell density (ECD), polymegathism, guttae density (GD) and guttae area ratio (GAR).

| Mouse ID | Age (months) | ECD (cells/mm2) | Polymegathism (%) | GD (guttae/mm2) | GAR (%) |
| --- | --- | --- | --- | --- | --- |
| 1 | 3 | 2306 | 25.23 | 40 | 0.70 |
| 2 | 3 | 2080 | 33.98 | 23 | 0.00 |
| 3 | 3 | 2658 | 22.99 | 0 | 0.58 |
| 4 | 3 | 2353 | 29.83 | 17 | 0.40 |
| 5 | 3 | 2533 | 24.87 | 12 | 1.55 |
| 6 | 3 | 2412 | 29.95 | 44 | 0.68 |
| 7 | 3 | 2320 | 28.46 | 23 | 1.96 |
| 8 | 3 | 1621 | 39.23 | 75 | 0.55 |
| 9 | 6 | 1448 | 55.63 | 73 | 2.70 |
| 10 | 6 | 1540 | 53.02 | 100 | 3.69 |
| 11 | 6 | 1510 | 52.50 | 71 | 4.27 |
| 12 | 6 | 1469 | 45.90 | 75 | 3.48 |
| 13 | 6 | 1680 | 50.46 | 115 | 3.96 |
| 14 | 6 | 1324 | 55.48 | 83 | 4.06 |
| 15 | 6 | 1558 | 40.16 | 80 | 2.93 |
| 16 | 9 | 1159 | 65.87 | 165 | 5.72 |
| 17 | 9 | 1217 | 55.19 | 109 | 3.96 |
| 18 | 9 | 1360 | 59.36 | 94 | 4.93 |
| 19 | 9 | 1371 | 45.93 | 50 | 2.12 |
| 20 | 9 | 1464 | 46.48 | 52 | 2.10 |
| 21 | 9 | 1264 | 56.57 | 60 | 4.79 |
| 22 | 9 | 1334 | 56.44 | 53 | 3.30 |
| 23 | 12 | 1180 | 61.46 | 128 | 2.24 |
| 24 | 12 | 1118 | 64.59 | 113 | 2.38 |
| 25 | 12 | 1167 | 61.71 | 149 | 2.85 |
| 26 | 12 | 943 | 82.25 | 226 | 4.65 |
| 27 | 12 | 1374 | 53.92 | 81 | 1.97 |
| 28 | 12 | 1229 | 56.09 | 153 | 2.73 |
| 29 | 12 | 1382 | 58.76 | 230 | 5.98 |
| 30 | 15 | 1178 | 62.52 | 140 | 4.01 |
| 31 | 15 | 1011 | 59.60 | 240 | 5.21 |
| 32 | 15 | 994 | 66.76 | 209 | 6.94 |
| 33 | 15 | 1085 | 56.28 | 201 | 4.54 |
| 34 | 15 | 990 | 66.20 | 241 | 5.42 |
| 35 | 15 | 1125 | 53.18 | 185 | 7.70 |
| 36 | 15 | 1121 | 61.18 | 187 | 2.30 |
| 37 | 18 | 1110 | 67.87 | 278 | 5.46 |
| 38 | 18 | 1198 | 64.24 | 182 | 4.89 |
| 39 | 18 | 1268 | 60.46 | 236 | 4.24 |
| 40 | 18 | 1293 | 75.59 | 160 | 5.59 |
| 41 | 18 | 1360 | 61.97 | 173 | 3.74 |
| 42 | 18 | 1147 | 71.21 | 230 | 3.83 |
| 43 | 18 | 1007 | 69.32 | 213 | 4.38 |
| 44 | 18 | 992 | 74.67 | 248 | 2.12 |
| 45 | 21 | 960 | 84.29 | 216 | 5.23 |
| 46 | 21 | 927 | 65.58 | 280 | 2.84 |
| 47 | 21 | 1075 | 66.93 | 291 | 4.86 |
| 48 | 21 | 1006 | 67.04 | 210 | 5.42 |
| 49 | 21 | 1228 | 70.13 | 232 | 3.74 |
| 50 | 21 | 963 | 72.99 | 265 | 4.71 |
| 51 | 21 | 1018 | 70.21 | 371 | 5.38 |
| 52 | 21 | 1176 | 60.92 | 307 | 11.24 |
| 53 | 24 | 1000 | 63.80 | 243 | 9.36 |
| 54 | 24 | 1085 | 59.15 | 239 | 5.35 |
| 55 | 24 | 972 | 66.75 | 332 | 6.86 |
| 56 | 24 | 1007 | 62.87 | 318 | 9.09 |
| 57 | 24 | 933 | 70.29 | 251 | 6.37 |
| 58 | 24 | 1073 | 60.80 | 254 | 6.26 |

**Supplementary Table 3.** Individual measurements for wildtype mice including age, endothelial cell density (ECD) and polymegathism.

| Mouse ID | Age (months) | ECD (cells/mm<sup>2</sup>) | Polymegathism (%) |
| --- | --- | --- | --- |
| 1 | 3 | 2440 | 17.56 |
| 2 | 3 | 2737 | 18.13 |
| 3 | 3 | 2648 | 17.05 |
| 4 | 3 | 2791 | 17.71 |
| 5 | 3 | 2841 | 14.42 |
| 6 | 3 | 2830 | 15.14 |
| 7 | 3 | 2713 | 16.91 |
| 8 | 6 | 2381 | 21.79 |
| 9 | 6 | 2475 | 17.81 |
| 10 | 6 | 2379 | 18.24 |
| 11 | 6 | 2480 | 17.44 |
| 12 | 6 | 2049 | 15.45 |
| 13 | 6 | 2309 | 15.41 |
| 14 | 6 | 2230 | 16.93 |
| 15 | 9 | 2162 | 23.33 |
| 16 | 9 | 2232 | 20.37 |
| 17 | 9 | 2443 | 16.51 |
| 18 | 9 | 2430 | 21.34 |
| 19 | 9 | 2409 | 20.64 |
| 20 | 9 | 2458 | 16.14 |
| 21 | 9 | 2284 | 16.46 |
| 22 | 12 | 2290 | 21.16 |
| 23 | 12 | 2252 | 16.99 |
| 24 | 12 | 2366 | 21.56 |
| 25 | 12 | 2131 | 21.41 |
| 26 | 12 | 2317 | 18.63 |
| 27 | 12 | 2259 | 16.87 |
| 28 | 12 | 2119 | 21.90 |
| 29 | 12 | 2437 | 20.01 |
| 30 | 15 | 2288 | 17.87 |
| 31 | 15 | 2212 | 20.26 |
| 32 | 15 | 2290 | 19.44 |
| 33 | 15 | 2378 | 17.95 |
| 34 | 15 | 2323 | 16.15 |
| 35 | 15 | 2306 | 19.62 |
| 36 | 15 | 2226 | 16.83 |
| 37 | 15 | 2060 | 15.88 |
| 38 | 18 | 2167 | 17.97 |
| 39 | 18 | 1970 | 21.30 |
| 40 | 18 | 2378 | 18.49 |
| 41 | 18 | 2433 | 21.74 |
| 42 | 18 | 2387 | 20.93 |
| 43 | 18 | 2393 | 18.87 |
| 44 | 18 | 2103 | 20.30 |
| 45 | 18 | 2623 | 23.49 |
| 46 | 21 | 2168 | 20.85 |
| 47 | 21 | 2143 | 24.20 |
| 48 | 21 | 2246 | 23.39 |
| 49 | 21 | 2129 | 18.32 |
| 50 | 21 | 2250 | 22.46 |
| 51 | 21 | 2123 | 21.72 |
| 52 | 21 | 2171 | 28.92 |
| 53 | 24 | 2282 | 21.77 |
| 54 | 24 | 2139 | 28.82 |
| 55 | 24 | 1877 | 33.73 |
| 56 | 24 | 2005 | 26.12 |
| 57 | 24 | 1933 | 30.54 |
| 58 | 24 | 2108 | 28.54 |

**Supplementary Table 4.** Descriptive demographic data for FECD patients that were used for the analysis of guttae anatomical location (vertices versus central endothelial cells) with specular microscopy (SPEC).

| **SPEC ID** | **Age**<br>**(years)** | **Sex** | **Krachmer**<br>**grade** | **Number of**<br>**images** |
| --- | --- | --- | --- | --- |
| SP091 | 76 | Female | 4 | 6 |
| SP093 | 70 | Male | 3 | 2 |
| SP096 | 74 | Female | 5 | 3 |
| SP114 | 63 | Male | 3 | 3 |
| SP115 | 84 | Male | 5 | 1 |
| SP117 | 67 | Female | 4 | 3 |
| SP118 | 90 | Female | 5 | 1 |
| SP001 | 76 | Female | 4 | 2 |
| SP004 | 70 | Female | 4 | 5 |
| SP006 | 50 | Female | 3 | 3 |
| SP007 | 59 | Female | 4 | 4 |
| SP020 | 81 | Female | 3 | 1 |
| SP028 | 80 | Female | 5 | 2 |
| SP028 | 80 | Female | 3 | 2 |
| SP030 | 77 | Female | 4 | 4 |
| SP008 | 47 | Male | 4 | 2 |
| SP010 | 76 | Female | 5 | 1 |
| SP010 | 76 | Female | 3 | 1 |
| SP013 | 76 | Male | 4 | 2 |
| SP014 | 81 | Male | 3 | 2 |

**Supplementary Table 5.** Descriptive demographic data for FECD patients that their corneas were used for the analysis of immunohistochemistry results.

| ID | Age<br>(years) | Sex | Medical History |
| --- | --- | --- | --- |
| P | 81 | Male | hypotension, peripheral vascular disease, cellulitis of right lower limb, Myasthenia gravis, foot surgery |
| Q | 65 | Female | Epiretinal membrane-OD , secondary IOL placement OD, pars plana vitrectomy OD, chalazion right lower lid, hysterectomy, hypertension, bulging discs, trigger finger release surgery |
| R | 72 | Male | Ocular hypertension, hyperlipidemia, migraines, tubular adenoma of colon, GED, coronary artery disease, hypercholesterolemia, CAD, previous cocaine use, Asthma, Colon polys, BPH, PND related AR & seasonal allergies, OSA w. APAP, ED, umbilical hernia, |
| S | 63 | Male | current tabacco use, Asthma, Diabetes type 2, hypertension, arthritis, prostate cancer, lung disease, hernia repair |
| T | 97 | Male | Hip joint replacement, osteoarthritis lower leg, blepharoplasty, hypertension, hyperlipidemia, sinus bradycardia, chronic dyspnea with obstructive lung disease, chronic macrocytosis, coronary artery disease, chronic kidney disease |
| 53 | 68 | Male | CKD stage 3 |
| 54 | 67 | Male | Hypertension, Asthma, Pigment dispersion syndrome, dry eyes, ocular hypertension-OU, appendectomy, hernia surgery, tonsillectomy, large benign tumor removed |
| 73 | 77 | Male | arthritis, hyperlipidemia, hypertension, GED, Rheumatoid arthritis, chronic systolic heart failure, squamous cell lung cancer, respiratory failure, St elevation myocardial infarction, previous tobacco use |
| 83 | 68 | Female | MGD, hypertensive retinopathy OU, previous tabacco use, high cholesterol, hypertension, asthma, dermatitis, murmur heart, thoracotomy, sleep apnea, depression-anxiety, fibromyalgia, acid reflux, total knee replacement-right, cartel tunnel |
| 101 | 71 | Female | Osteoporosis, allergic rhinitis, atrophic vaginitis, Dyspareunia, GERD, Essential Hypertension, Conjunctivoplasty OD, dietary calcium deficiency, tubular adenoma of colon, diverticulosis of colon, reactive hypoglycemia, |
| 142 | 72 | Male | Bilateral Glaucoma, Chronic iritis OS-HZO, trabeculectomy OS, Hypertension, paroxysmal atrial fibrillation, coronary artery disease, pulmonary embolism, stent of left PDA, rotator cuff arthropathy of both shoulders |
| 220 | 84 | Female | Atrial fibrillation, hypothyroidism, GERD, OSA, Diastolic dysfunction, neurocognitive disorder, HSV, mixed hyperlipidemia, chronic diastolic heart failure, Cervical Cancer, blepharitis OU |
| 276 | 47 | Female | Epilepsy, thyroid cyst removed from neck, Grand Mal Seizure, right & left Lung partially deflated, eczema, previous tobacco use including vaping |
| 277 | 83 | Female | Optic Cupping-OU, Dry Eye Syndrome, Brian conditions NEC, Rathke cleft cyst s/p resection, Hematometra, cholecystectomy, hysterectomy, blepharoplasty, septoplasty, transsphenoidal hypophysectomy/ resection pituitary tumor |
| 278 | 71 | Male | cystoid macular edema OD, dyslipidemia, benign prostatic hyperplasia with lower urinary tract symptoms, appendectomy, sinus surgery, toe surgery, neck surgery |
| 280 | 69 | Female | hypothyroidism, hyperlipidemia, GERD, Barrett's esophagus, sleep apnea, factor V Leiden, thyroid nodule, blepharochalasis, appendectomy, hysterectomy, hernia, abdomen surgery, Osteoarthritis, cardiac radiofrequency ablation, hiatal hernia, premature ventricular contraction |
| 281 | 68 | Male | hypothyroidism, hyperlipidemia, GERD, Barrett's esophagus, sleep apnea, factor V Leiden, thyroid nodule, blepharochalasis, appendectomy, hysterectomy, hernia, abdomen surgery, Osteoarthritis, cardiac radiofrequency ablation, hiatal hernia, premature ventricular contraction |

**Supplementary Table 6.** Descriptive demographic data for donor patients that their corneas were used for the analysis of immunohistochemistry results.

| ID | Age<br>(years) | Sex | Medical History |
| --- | --- | --- | --- |
| 22-001085 | 62 | Male | Hyperlipidemia, cocaine use, previous tobacco use, ethanol use, LASIK (bilateral). |
| 22-001113 | 76 | Male | Coronary artery disease, hematuria, hyperlipidemia, a-fib, gastrointestinal bleeding, anemia, asthma, cerebellar infarction, chronic heart failure, fluid retention, previous tobacco use, ethanol use, osteoporosis, urinary retention, renal failure, dilated cardiomyopathy, clostridioides difficile, urinary tract infection, fever, cholecystectomy, hernia repair, wisdom teeth. |
| 25-001395 | 69 | Male | Short of breath, bronchitis, chronic cough, COPD, emphysema, allergies, tobacco use, ethanol use, rheumatoid arthritis, ST-segment elevation myocardial infarction, left shoulder surgery, bilateral knee surgery. |
| 24-000625 | 60 | Male | Iron deficiency, blood loss, subclinical hypothyroidism, tobacco use, recent syncope, hypertension, allergic to augmentin, marijuana use, ethanol use, colonoscopy, mouth surgery, esophagogastroduodenoscopy. |
| 24-000669 | 65 | Male | Secondary polycythemia, morbid obesity, rib fracture, transverse process lumbar fracture, glaucoma, dry eye syndrome, right lower extremity lymphedema, hyperlipidemia, left carotid artery stenosis, ethanol abuse, hypertension, peripheral vascular disease, 100% occlusion of left carotid artery, stroke with right lateral Webril hemisphere necrosis, seizure disorder, hematoma of scalp, lamina papyracea fracture, emphysema, myocardial infarction, swelling with discoloration of bilateral lower extremities, chronic cough, diarrhea, blood blisters and sores on hands and wrists, seizure, tremors of hands/arms, difficulty walking, confusion, marijuana use, tobacco use, heart valve procedure. |
| 24-000697 | 74 | Female | Glioblastoma multiforme of the parietal lobe (grade 4), chemotherapy, radiation therapy, hypokalemia, left homonymous hemianopia, vasogenic cerebral edema, bilateral glaucoma suspect, malignant neoplasm of brain, thrombocytopenia, iron deficiency anemia, visual disturbance, weakness, weight loss, cancer rash of chest arms, and legs, mental decline, hallucinations, difficulty walking, former tobacco use, ethanol use, tubal ligation, craniotomy, visula disturbance, left homonymous hemianopia. |
| 24-000702 | 42 | Male | Foot fracture, allergic to penicillin, tobacco use, ethanol use, kidney stone, eye sty. |
| 25-001405 | 60 | Female | Anemia, low white blood cell counts, idiopathic aplastic anemia, athritis, hypertension, mental disorder, migraine, asthma, heart disease, autoimmune disease, transient ischemic attack, visual disturbances, cerebral cavernoma, visual disturbances. |
| 25-11-034 | 66 | Male | Coronary artery disease, hypertension, high cholesterol, 6x vascular stents. |
| 25-001205 | 65 | Male | Cholangiocarcinoma, radiation, oral chemotherapy, bile duct cancer, coronary artery disease, irritable bowel syndrome, hypertension, recurrent stroke, hyperlipidemia, chronic heart failure, bilateral vestibular hypofunction, deep vein thrombosis, pulmonary embolism, chronic anemia, adjustment disorder, malignant neoplasm metastatic to lung, metastasis to peritoneum, appendectomy, cholecystectomy, small bowel surgery, coronary artery bypass grafting, tonsillectomy, urethral stent, peripheral vision loss, confusion, headache, peripheral vision loss. |
| 25-001264 | 78 | Male | Aortic aneurysm, dialysis, oxygen, hand tremors, difficulty walking, weakness, pacemaker, former tobacco use, ethanol use, hypertension, hyperlipidemia, heart attack, black esophagus, diverticulitis, abdominal aortic aneurysm, dysphagia, acute hypoxic respiratory failure, esophageal necrosis, acute oliguric renal failure, COPD, a-fib, obstructive sleep apnea, peripheral vascular disease, acute kidney injury, ventricular tachycardia, esophageal obstruction, acute hypoxic respiratory failure, epistaxis, multifocal pneumonia, bilateral pleural effusion, anuria, acute blood loss anemia, mood disorder, acute tubular necrosis, hemodialysis, ischemic cardiomyopathy, heart failure with preserved ejection fraction, coronary artery disease, aneurysm surgery, bilateral knee replacements, pacemaker, triple bypass, hemodialysis, coronary artery bypass grafting, esophagogastroduodenoscopy. |
| 25-001287 | 70 | Female | Breast cancer, COPD, chemotherapy, radiation therapy, tobacco use, a-fib, chronic heart failure, coronary artery disease, cubital tunnel syndrome, depression, gout, IgG deficiency, memory difficulty, skin cancer, ethanol use, cyst, hand tingling/numbness, acute exacerbation of chronic obstructive airway disease, cellulitis, hypoxia, bresat lumpectomy, cardiac stents, nasal skin graft, cyst removal, tonsillectomy, tubal ligation, surgical removal of palate, alveoplasty, teeth extraction. |
| 25-001201 | 71 | Male | Asthma, ethanol use, hypertension, hyperlipidemia, varicella, appendectomy, colonoscopy with polypectomy. |

